# Identification of two glycosyltransferases required for synthesis of membrane glycolipids in *Clostridioides difficile*

**DOI:** 10.1101/2025.01.14.632984

**Authors:** Brianne R. Zbylicki, Sierra Cochran, David S. Weiss, Craig D. Ellermeier

## Abstract

*Clostridioides difficile* infections cause over 12,000 deaths and an estimated one billion dollars in healthcare costs annually in the United States. The cell membrane is an essential structure that is important for protection from the extracellular environment, signal transduction, and transport of nutrients. The polar membrane lipids of *C. difficile* are ∼50% glycolipids, a higher percentage than most other organisms. The glycolipids of *C. difficile* consist of monohexosyldiradylglycerol (MHDRG) (∼14%), dihexosyldiradylglycerol (DHDRG) (∼15%), trihexosyldiradylglycerol (THDRG) (∼5%), and a unique glycolipid aminohexosyl-hexosyldiradylglycerol (HNHDRG) (∼16%). Previously, we found HexSDF are required for synthesis of HNHDRG. The enzymes required for synthesis of MHDRG, DHDRG, and THDRG are not known. In this study, we identified the glycosyltransferases UgtA (CDR20291_0008), which is required for synthesis of all glycolipids, and UgtB (CDR20291_1186), which is required for synthesis of DHDRG and THDRG. We propose a model where UgtA synthesizes only MHDRG, HexSDF synthesize HNHDRG from MHDRG, and UgtB synthesizes DHDRG and potentially THDRG from MHDRG. We also report that glycolipids are important for critical cell functions, including sporulation, cell size and morphology, maintaining membrane fluidity, colony morphology, and resistance to some membrane targeting antimicrobials.

**Importance:** *Clostridioides difficile* infections are the leading cause of healthcare associated diarrhea. *C. difficile* poses a risk to public health due to its ability to form spores and cause recurrent infections. Glycolipids make up ∼50% of the polar lipids in the *C. difficile* membrane, a higher percentage than other common pathogens and include a unique glycolipid not present in other organisms. Here, we identify glycosyltransferases required for synthesis of glycolipids in *C. difficile* and demonstrate the important role glycolipids play in *C. difficile* physiology.

## Introduction

*Clostridioides difficile* is a Gram-positive, spore-forming, obligate anaerobe, and an opportunistic pathogen the Centers for Disease Control (CDC) has classified as an urgent threat to public health. According to the CDC, *C. difficile* infections caused over 12,000 deaths and an estimated one billion dollars in healthcare costs in 2019 in the United States ^1^. *C. difficile* infections are common in people who have undergone antibiotic treatment and can cause symptoms ranging from mild self-limiting diarrhea to life threatening pseudomembranous colitis ^2,3^. *C. difficile* infections are among the most common causes of healthcare-associated diarrhea ^4^.

*C. difficile* produces metabolically dormant spores that can persist aerobically and can resist antibiotic treatment and many disinfectants ^5–7^. Spores allow *C. difficile* to be transmitted aerobically and cause recurrent infections ^8^. To resume vegetative growth, spores require germinants and co-germinants ^9^. The primary germinant for *C. difficile* spores is the conjugated primary bile acid taurocholate (TCA), but other bile acids like glycocholate (GCA), deoxycholate (DCA), and cholate (CA) can also induce germination^9^. The co-germinant is often an amino acid like glycine or L-alanine ^9,10^. The primary bile acids CA and chenodeoxycholate (CDCA) are produced in the liver from cholesterol and can be conjugated in the liver with taurine or glycine ^11^. Once in the intestinal tract, the conjugated primary bile acids undergo processing via the intestinal microbiota and are de-conjugated by bile salt hydrolases and further dehydroxylated to make secondary bile acids ^11^. The secondary bile acids lithocholate (LCA) and DCA inhibit *C. difficile* vegetative cells after germination ^9,12,13^.

As a Gram-positive organism, *C. difficile* is protected from its extracellular environment by its cell envelope which includes a proteinaceous S-layer, a thick layer of peptidoglycan, cell envelope associated polysaccharides, and a cell membrane. These cell wall-associated polysaccharides include wall teichoic acids (WTA) and lipoteichoic acids (LTA), also called PS-II and PS-III in *C. difficile*, respectively ^14,15^. The LTA is anchored in the membrane by triglucosyldiacylglyerol ^15^. The cell membrane is essential and its composition in *C. difficile* is distinct from model organisms. In model organisms such as *Escherichia coli* and *Bacillus subtilis*, the lipids that make up the membrane and their synthesis have been well defined. In *E. coli*, the inner membrane is composed of ∼75% phosphatidylethanolamine, ∼20% phosphatidylglycerol, and ∼5% cardiolipin and in *B. subtilis*, the membrane is composed of ∼25% phosphatidylethanolamine, ∼40% phosphatidylglycerol, ∼15% lysyl-phosphatidylglycerol, ∼2% cardiolipin, and ∼10% glycolipids ^16–19^. The composition of the polar membrane lipids of *C. difficile* 630 was determined to be ∼30% phosphatidylglycerol, ∼16% cardiolipin, and ∼50% glycolipids ^20^. Notably, *C. difficile* only contains the phospholipids phosphatidylglycerol and cardiolipin and is lacking phosphatidylethanolamine and phosphatidylserine that are common in other bacteria ^20^. The synthesis of phosphatidylglycerol in *C. difficile* is an essential process, and when the enzymes for phosphatidylglycerol synthesis (CdsA and PgsA) are knocked down using CRISPR interference (CRISPRi ), the cells have severe viability and morphological defects ^21^. In *C. difficile,* ClsA and ClsB synthesize cardiolipin from phosphatidylglycerol ^21^. In contrast little is known about glycolipid synthesis in *C. difficile*.

In *C. difficile,* the glycolipids make up ∼50% of the polar lipids in the membrane, a very high percentage compared to ∼11% glycolipids in *B. subtilis* and ∼9% in *Staphylococcus aureus* ^17–20,22^. The glycolipids of *C. difficile* consist of monohexosyldiradylglycerol (MHDRG) (∼14% of total membrane lipids), dihexosyldiradylglycerol (DHDRG) (∼15%), trihexosyldiradylglycerol (THDRG) (∼5%), and aminohexosyl-hexosyldiradylglycerol (HNHDRG) (∼16%) ^20^. HNHDRG is a novel amino-glycolipid that is only found in *C. difficile.* Previously we demonstrated that HexSDF are required for synthesis of HNHDRG and that loss of HNHDRG decreases daptomycin and bacitracin resistance^23^. Loss of HexSDF did not alter the synthesis of the other glycolipids, and the genes required for their synthesis are not known ^23^.

Of note, glycolipids in other organisms commonly contain glucose or galactose, but the specific sugars in *C. difficile* glycolipids are unknown, so they are currently called hexosyl. The lipids of *C. difficile* are found in both the more familiar diacyl form and the less common plasmalogen form in which one of the two fatty acids is joined to the glycerol by an ether instead of an ester linkage ^20,24^. *C. difficile* glycolipids have both diacyl and plasmalogen forms, thus we refer to them broadly as diradylglycerols ^20^.

Some organisms have a single glycosyltransferase that processively synthesizes mono-, di- and tri-glycolipids, while other organisms utilize multiple enzymes, and the reason for this difference is currently unknown. *B. subtilis* and *S. aureus* have a single glycolipid glycosyltransferase (UgtP and YpfP respectively) that processively synthesizes monoglucosyldiacylglycerol (Glc-DAG) and diglucosyldiacylglycerol (Glc_2_-DAG) glycolipids ^25–27^. However, some organisms such as *Enterococcus faecalis* and *Streptococcus agalactiae* utilize two different glycosyltransferases to synthesize glycolipids. BgsB and BgsA from *E. faecalis* and Gbs0683 and IagA from *S. agalactiae* act sequentially to synthesize Glc-DAG and then Glc_2_-DAG in their respective organisms ^25,28–31^. Other organisms such as *Listeria monocytogenes* and *Streptococcus pneumoniae* first utilize a glycosyltransferase (LafA and Spr0982 respectively) to synthesize Glc-DAG, which is then used as a substrate for a second glycosyltransferase (LafB and CpoA respectively) to synthesize galactosylglucosyldiacylglycerol (GalGlcDAG) ^25,32–34^.

The enzymes required for synthesis of the glycolipids MHDRG, DHDRG, and THDRG in *C. difficile* are not known. Here we report the identification of two enzymes required for glycolipid synthesis in *C. difficile*; UgtA, a glycosyltransferase required for all glycolipid synthesis, and UgtB, a glycosyltransferase required for the synthesis of DHDRG and THDRG. We demonstrate that loss of all glycolipids in *C. difficile* reduces growth, alters colony and cell morphology, decreases sporulation frequency, decreases resistance to a subset of membrane-targeting antimicrobials, and increases membrane fluidity. We propose a model for glycolipid synthesis that starts with synthesis of MHDRG by UgtA (Fig. 1). MHDRG is then used by either HexSDF to synthesize HNHDRG or by UgtB to synthesize DHDRG and possibly THDRG from MHDRG.

**Figure 1.**
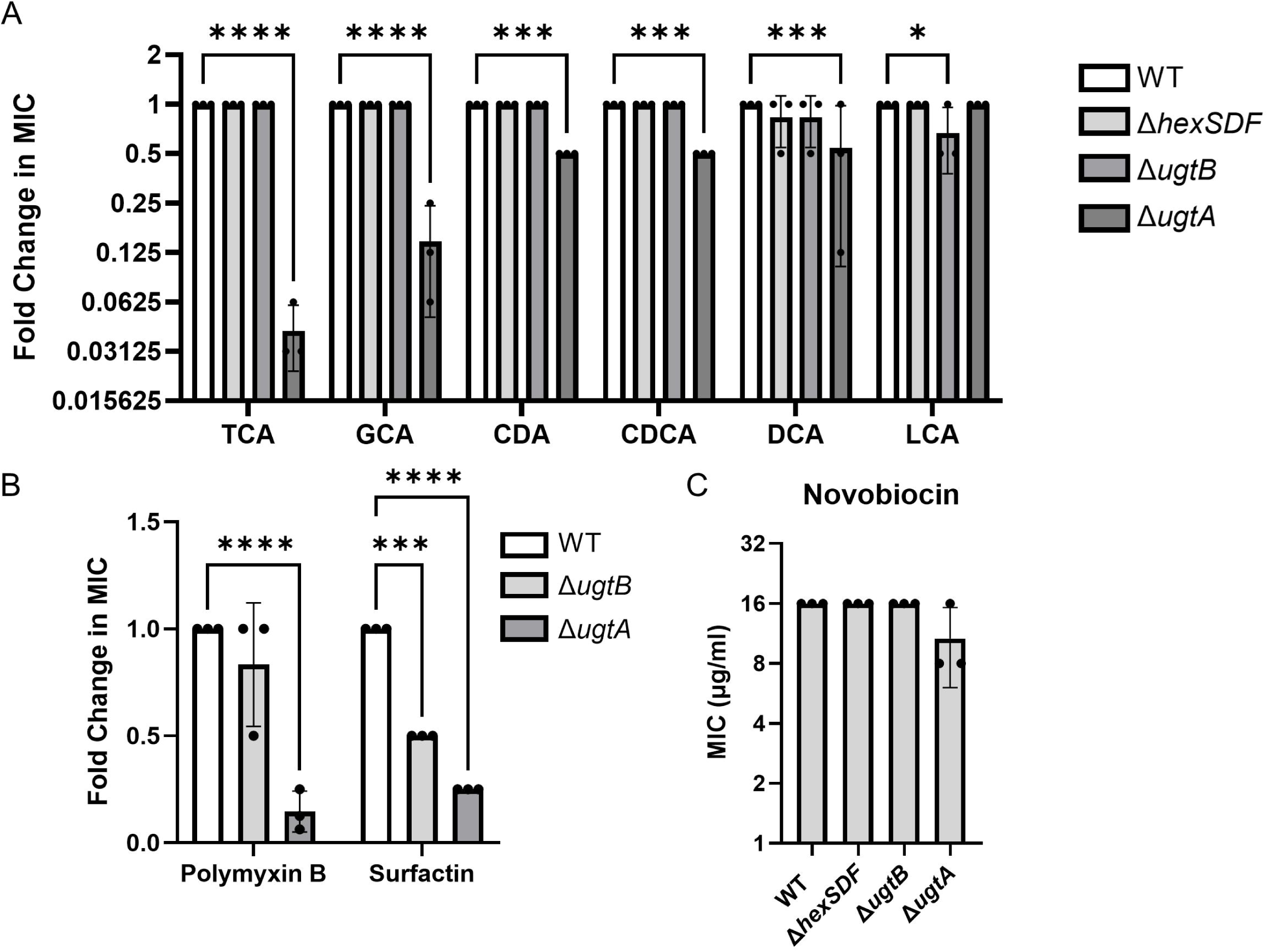
Model of glycolipid synthesis in *C. difficile*. UgtA synthesizes MHDRG, the precursor for all other glycolipids. UgtB synthesizes DHDRG using MHDRG as a substrate. UgtB may processively synthesize THDRG from DHDRG. We hypothesize that HexS adds *N*-acetyl-hexose to MHDRG to make a HNHDRG intermediate HNacHDRG, which then gets flipped to the outside of the cell by HexF or other flippases, and finally deacetylated by HexD to form HNHDRG. The localization of the glycolipids has not been experimentally determined, and if MHDRG, DHDRG, and THDRG exist on the outer leaflet of the membrane, the flippases involved are currently unknown.

## Results

### Identifying putative glycosyltransferases

HexSDF are required for synthesis of HNHDRG ^23^. However, the enzymes responsible for synthesis of MHDRG, DHDRG, and THDRG are not known. We performed a BLASTP search against *C. difficile* R20291 using BioCyc and sequences of glycosyltransferases from organisms which utilize processive glycosyltransferases *B. subtilis* (UgtP) and *S. aureus* (YpfP) or organisms which use multiple glycosyltransferases *E. faecalis* (BgsA and BgsB), *L. monocytogenes* (LafA and LafB), *S. agalactiae* (IagA and GBS0683) and *S. pneumoniae* (CpoA and SPR0982) ^35,36^. We identified a subset of 7 putative glycosyltransferases that met a threshold p-value of <1×10^−5^ to at least one of the proteins used to query the database (Table S1). An alignment made using Clustal Omega of the seven putative glycosyltransferases identified in *C. difficile* along with the glycosyltransferases from other organisms is shown in Fig. S1 ^37^. We chose *cdr20291_0008* (named *ugtA*), *cdr20291_1186* (named *ugtB*), *cdr20291_0773*, and *cdr20291_2958* for further study. We chose not to follow up on *cdr20291_2539* (*murG*)*, cdr20291_2614* (*hexS*), and *cdr20291_2658* because these already have known functions in synthesis of lipid II, HNHDRG and PS-II, respectively^23,38,39^.

### UgtA and UgtB are required for glycolipid synthesis

We used CRISPR mutagenesis to construct Δ*ugtA,* Δ*ugtB,* Δ*cdr0773,* and Δ*cdr2958* mutants. We grew strains to mid-log phase and then extracted lipids as previously described ^40^. We separated lipids using TLC and stained for glycolipids using 1-naphthol ^40^. We found the Δ*ugtA* mutant lacks all glycolipids and the Δ*ugtB* mutant lacks DHDRG and THDRG but still produces MHDRG and HNHDRG (Fig. 2A). Deletion of *cdr0773* or *cdr2958* did not alter the glycolipid profile. To ensure that *cdr0773* and *cdr2958* were not redundant, we constructed and tested a Δ*cdr0773* Δ*cdr2958* mutant and did not observe any change in the glycolipid profile compared to wild type (Fig. 2A).

**Figure 2.**
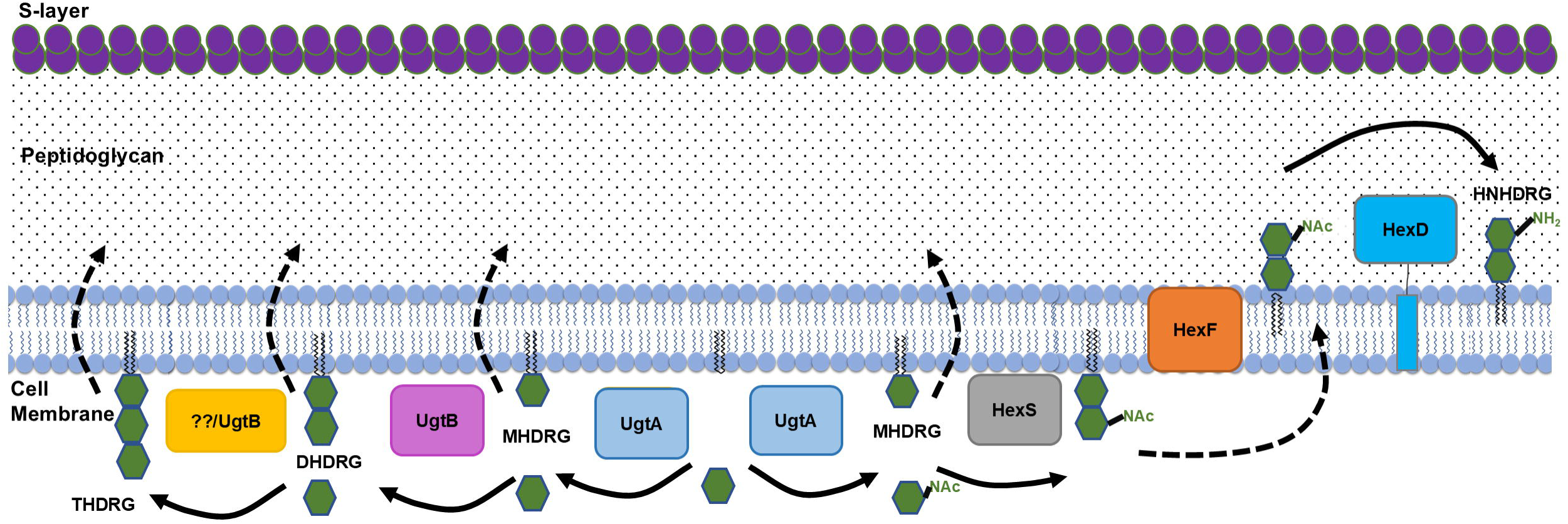
UgtA is required for glycolipid synthesis and UgtB is required for DHDRG and THDRG synthesis. A) Lipid extracts from wild type (WT), Δ*hexSDF,* Δ*ugtB,* Δ*ugtA,* Δ*cdr0773,* Δ*cdr2958,* and Δ*cdr0773* Δ*cdr2958* were separated using TLC and visualized with 1-naphthol. B) Lipid extracts from WT with an empty vector pAP114 (EV), Δ*ugtB* EV, Δ*ugtB* P*_xyl_*-*ugtB,* Δ*ugtA* EV, and Δ*ugtA* P*_xyl_*-*ugtA* were separated by TLC and visualized with 1-naphthol. Lipid purification and TLC was performed at least three separate times, and a representative example is shown. Comparison of C) MHDRG, D) DHDRG, E) HNHDRG, and F) THDRG levels as determined by lipidomic analysis for WT, Δ*ugtB*, and Δ*ugtA*. Strains carrying a plasmid were induced with 1% xylose to express elements of interest. Data are graphed as the mean and standard deviation of three replicates. Statistical significance was assessed by one-way analysis of variance using Dunnett’s multiple-comparison test. **** *p* < 0.0001, *** *p* < 0.001, ** *p* < 0.01, * *p* < 0.05.

We performed lipidomic analysis on the Δ*ugtA* and Δ*ugtB* mutants to quantify the loss of some or all of the glycolipids. The Δ*ugtA* mutant lacked all glycolipids and the Δ*ugtB* mutant lacked DHDRG and THDRG, corroborating the TLC data (Fig. 2C-F). To confirm that the loss of glycolipids was due to the loss of either *ugtA* or *ugtB,* we expressed *ugtA* or *ugtB* from a xylose-inducible promoter on a plasmid in the Δ*ugtA* and Δ*ugtB* mutants, respectively. We saw restoration of the respective missing glycolipids with both TLC and lipidomic analysis (Fig. 2B, S2). There was no obvious change in the phospholipids phosphatidylglycerol or cardiolipin with loss of either *ugtA* or *ugtB* (Fig. S2). Our data suggest UgtA is required for synthesis of MHDRG and UgtB is required for DHDRG and THDRG.

### UgtA and UgtB are sufficient to synthesize glycolipids

We hypothesized that UgtA produces MHDRG, which can serve as a substrate for either UgtB to produce DHDRG or HexSDF to produce HNHDRG (Fig. 1). To test this idea, we sought to determine if UgtA or UgtB is sufficient to synthesize glycolipids in a heterologous host. In *B. subtilis* UgtP is required for synthesis of all glycolipids ^26^. Thus, we expressed either *ugtA* or *ugtB* in a *B. subtilis* Δ*ugtP* mutant and examined the glycolipid profiles using TLC and lipidomic analysis. When *ugtA* was expressed alone, MHDRG was produced, demonstrating that *ugtA* is sufficient to synthesize MHDRG (Fig. 3A, B). When *ugtB* was expressed alone, no glycolipids were detected (Fig. 3). When *ugtA* and *ugtB* were expressed together, MHDRG was detected but we did not detect DHDRG by TLC, however a small amount of DHDRG was detected via lipidomic analysis (Fig. 3A, C). These findings suggest that UgtB utilizes MHDRG as a substrate to produce DHDRG.

**Figure 3.**
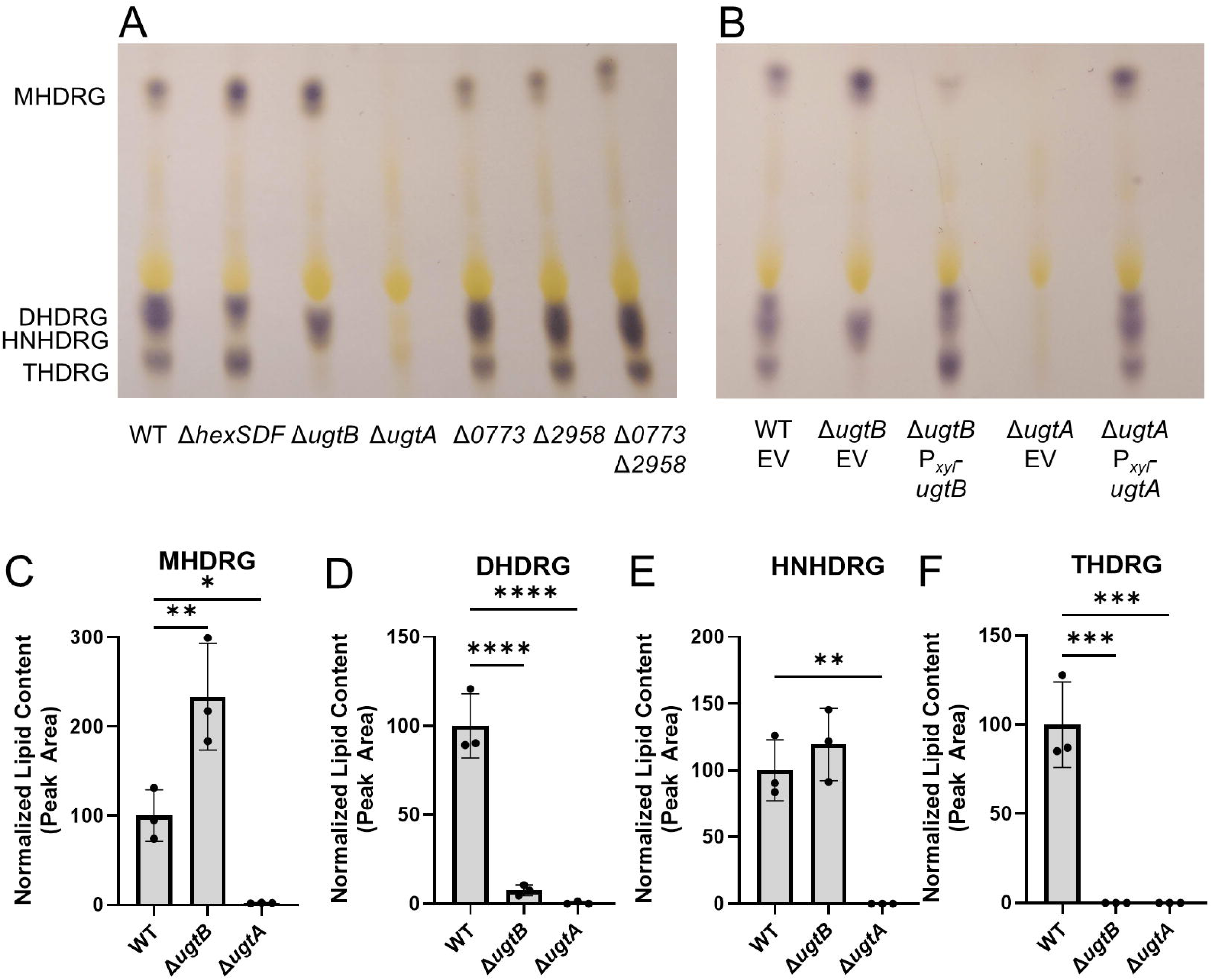
Exogenous expression of *ugtA*, *ugtB,* and *hexSDF* in *B. subtilis* supports production of glycolipids. A) Lipid extracts from *B. subtilis* strains were separated using TLC and visualized with 1-naphthol. Comparison of B) MHDRG, C) DHDRG, and D) HNHDRG levels as determined by lipidomic analysis. The black arrow highlights the small amount of DHDRG in Δ*ugtP* P*_xyl_*-*ugtA* P*_IPTG_*-*ugtB.* Strains were induced with 1% xylose and/or 1 mM IPTG to express elements of interest.

### HexSDF are sufficient to synthesize HNHDRG from MHDRG

We previously showed that HexSDF were required for synthesis of the unique glycolipid HNHDRG, and we proposed that HNHDRG is synthesized from MHDRG (Fig. 1) ^23^. To test this, we expressed *hexSDF* in a Δ*ugtP* mutant of *B. subtilis.* When *hexSDF* were expressed alone, no glycolipid, including HNHDRG, could be detected via TLC or lipidomic analysis (Fig. 3A, D). However, when *hexSDF* were co-expressed with *ugtA* in a Δ*ugtP* mutant, both MHDRG and HNHDRG are produced and can be observed by TLC and lipidomic analysis (Fig. 3A, D). This supports the model that UgtA produces MHDRG and HexSDF synthesizes HNHDRG using MHDRG as a substrate (Fig. 1).

### Loss of UgtA alters growth and colony morphology

We sought to determine the effects of loss of different glycolipids on the physiology of *C. difficile*. The Δ*ugtA* mutant reached the same OD_600_ over 24 hours but had a modest yet clear growth defect when compared to wild type (Fig. 4A, S3). Expression of *ugtA* in *trans* in the Δ*ugtA* mutant restored wild-type growth rates (Fig. 4A). In contrast neither the Δ*ugtB* nor Δ*hexSDF* mutants showed altered growth rates (Fig. S3). We plated 10-fold dilutions of an overnight culture onto a TY plate and imaged the resulting colonies after 24 hours. The Δ*ugtA* mutant grew to the same spot dilution as the wild-type control, suggesting there is not a strong viability defect, but the colonies were smaller and had a smoother morphology (Fig. 4B, S3B). The colony morphology of Δ*ugtA* was restored to wild type when UgtA was produced from a plasmid (Fig. 4B). The Δ*ugtB* and Δ*hexSDF* mutants had similar growth and colony morphology as wild type (Fig. S3).

**Figure 4.**
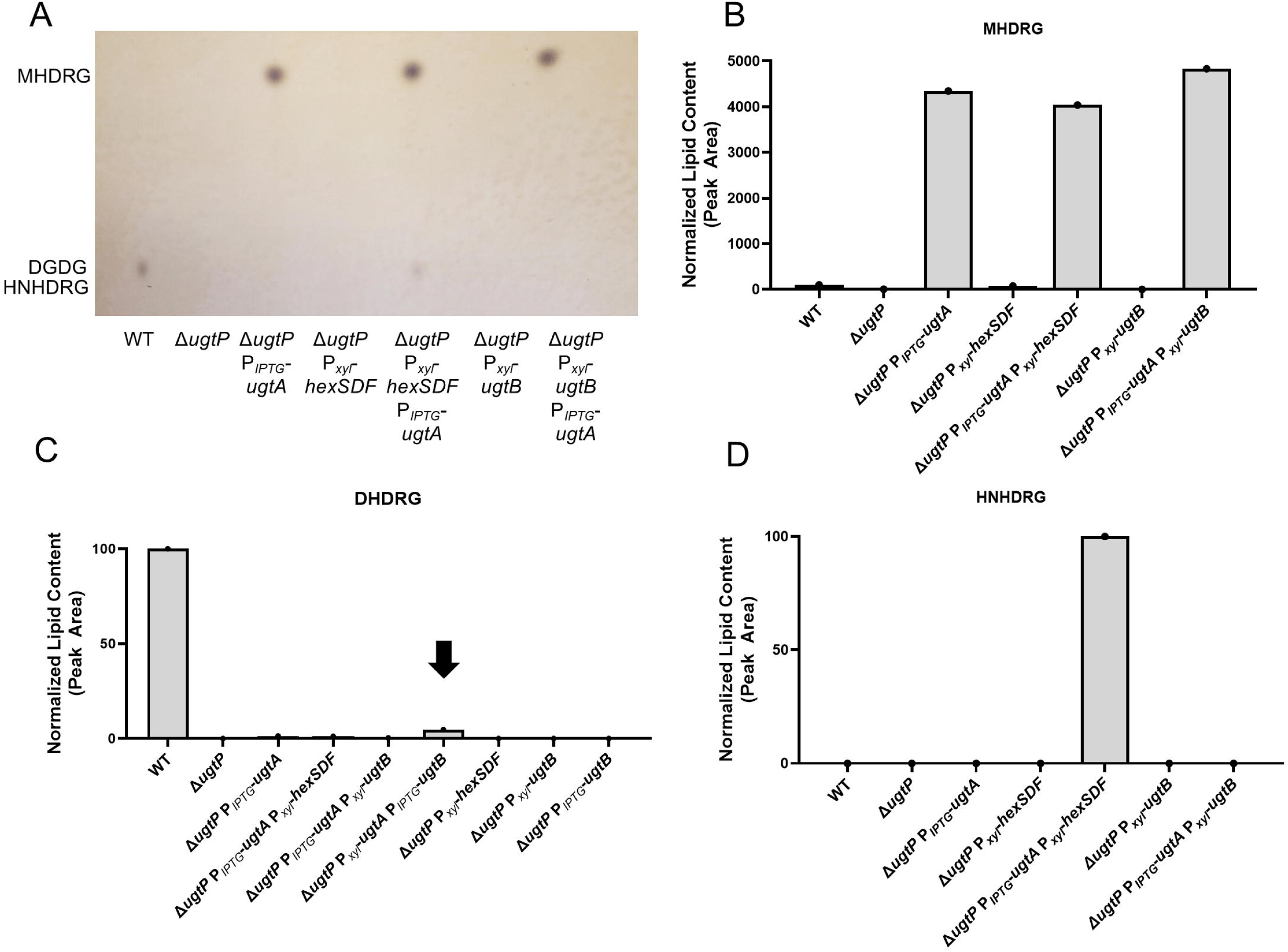
Loss of glycolipids alters growth and cell and colony morphology. A) Overnight cultures of WT EV, Δ*ugtA* EV, and Δ*ugtA* P*_xyl_*-*ugtA* were subcultured into TY Thi10 1% xylose to an OD_600_ of 0.05 and growth was measured using OD_600_. Data are graphed as the mean and standard deviation of three replicates. B) 10-fold dilutions of overnight cultures of WT EV, Δ*ugtA* EV, and Δ*ugtA* P*_xyl_*-*ugtA* grown in TY Thi10 were plated onto TY Thi10 1% xylose and the resulting colonies were imaged after 24 hours. C) Phase contrast microscopy and D) fluorescence microscopy using membrane stain FM4-64 of WT, Δ*hexSDF,* Δ*ugtB,* and Δ*ugtA*. Images shown are representative of three independent experiments. E) Cell length as calculated from at least 300 cells from three independent experiments. Dots represent individual cells and are colored red, yellow or blue to distinguish the three experiments. The mean of each experiment is indicated by a black circle, square or triangle. The horizontal bar and whiskers depict the mean and standard deviation of the three experiments. F) Septa/cell as calculated from at least 300 cells from three independent experiments. Data are graphed as the mean and standard deviation, with inverted triangles for individual cells. Percent of cells with >1 septa/cell are indicated. Statistical significance was assessed by one-way analysis of variance using Dunnett’s multiple-comparison test. **** *p* < 0.0001.

### The Δ*ugtB* and Δ*ugtA* mutants have altered cell morphology

Since *ugtA* mutants have altered growth rates we sought to determine the effect of loss of glycolipids on cell morphology. We found that Δ*ugtA* mutant cells are the same length as wild type, but they are curvier as measured by sinuosity and have more septa per cell as revealed by staining with the membrane dye FM4-64 (Fig. 4C-F, S4A,C). In contrast, deletion of *ugtB* resulted in a subtle increase in cell length that was not statistically significant (Fig. 4C-E). We complemented the Δ*ugtA* deletion by expressing *ugtA* in *trans* and observed cell sinuosity and septa per cell restored to wild-type levels (Fig. S4A,B,E,F). However, in the complementation experiments, the phenotypes exhibited by Δ*ugtA* and Δ*ugtB* with an empty vector control are not as severe as the deletions alone. We hypothesize that this is due to decreased growth rates when cells are grown under antibiotic pressure to maintain plasmids.

In *B. subtilis*, a Δ*ugtP* mutant has rough, aberrant structures on the cell surface ^41^. We examined the Δ*ugtA,* Δ*ugtB,* and Δ*hexSDF* mutants carrying an empty vector using TEM and did not see any note worthy changes in the cell surface structure compared to the wild-type control (Fig. S4).

### Loss of UgtA decreases sporulation

Glycolipids produced by the glycosyltransferase UgtP are present in spores of *B. subtilis* ^18,19^. While the lipid composition of *C. difficile* spores has not been determined, we wanted to know if the loss of various glycolipids affected sporulation. We tested the sporulation frequencies of wild type, Δ*ugtA,* Δ*ugtB*, and Δ*hexSDF* mutants as previously described ^42^. We found the Δ*ugtA* mutant has a ∼25-fold decrease in sporulation frequency compared to wild type, and sporulation could be rescued by expression of *ugtA* from a plasmid (Fig. 5A, S3F). The sporulation frequencies of Δ*hexSDF* and Δ*ugtB* mutants were similar to wild type (Fig. 5A).

**Figure 5.**
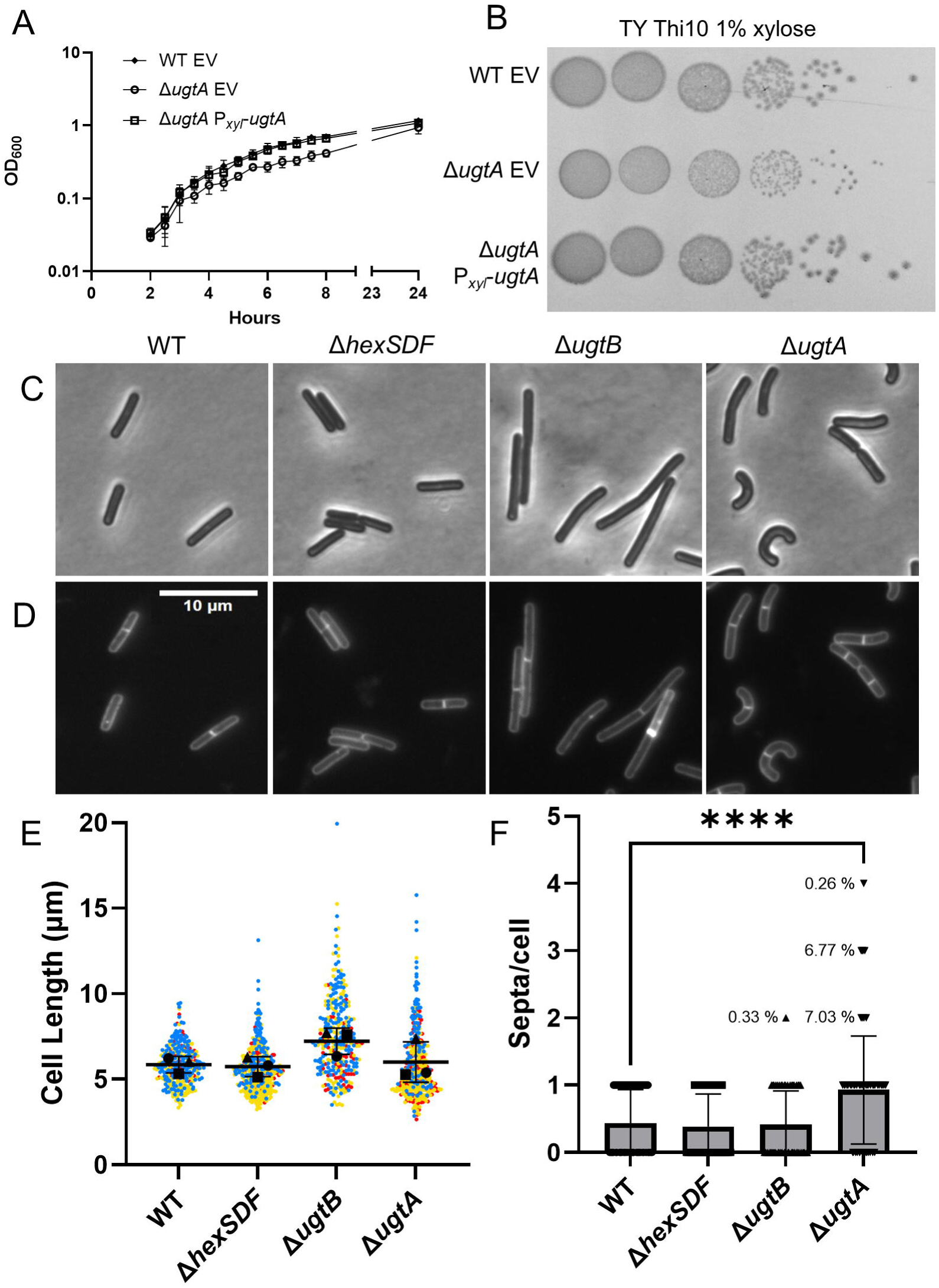
Loss of glycolipids decreases sporulation frequency, alters colony morphology in the presence of taurocholate, and increases membrane fluidity. A) Sporulation frequency of WT, Δ*hexSDF,* Δ*ugtB,* and Δ*ugtA.* 10-fold dilutions of overnight cultures of WT EV, Δ*ugtA* EV, and Δ*ugtA* P*_xyl_*-*ugtA* grown in BHIS Thi10 were plated on B) BHIS Thi10 1% xylose and C) BHIS Thi10 1% xylose 0.1% TCA. D) Relative membrane fluidity of WT EV, Δ*ugtA* EV, and Δ*ugtA* P*_xyl_*-*ugtA* normalized to WT EV. Strains carrying a plasmid were induced with 1% xylose to express elements of interest. Data are graphed as the mean and standard deviation from three replicates. Statistical significance was assessed by one-way analysis of variance using Dunnett’s multiple-comparison test. ** *p* < 0.01, * *p* < 0.05.

### Loss of UgtA increases sensitivity to some bile acids

Upon plating Δ*ugtA* spores on germination media (BHIS 0.1% TCA) the resulting colonies were very small. To investigate this further we plated vegetative cells from wild type, Δ*ugtA,* Δ*ugtB,* and Δ*hexSDF* on TY, BHIS, and BHIS + 0.1% TCA. We found that Δ*ugtA* produces small colonies on TY and BHIS and this small colony phenotype was exacerbated on BHIS 0.1% TCA (Fig. 5B-C, S3B,D,E). In contrast deletion of *hexSDF* or *ugtB* did not alter colony morphology when plated on BHIS 0.1% TCA (Fig. S3E). The expression of *ugtA* in the Δ*ugtA* mutant restored colony morphology to wild-type under all conditions (Fig. 4B, 5B-C).

Because of these observations we then tested the sensitivity of wild type, Δ*ugtA,* Δ*ugtB,* and Δ*hexSDF* to several bile acids including TCA, GCA, CA, CDCA, DCA, and LCA using minimum inhibitory concentrations assays (MICs). We found that the Δ*ugtA* mutant has a ∼25-fold decrease in TCA and a >7-fold decrease in GCA resistance compared to wild type (Fig. 6A). These defects could be complemented by expressing *ugtA* in *trans* (Fig. S5). The Δ*ugtA* mutant was also slightly more sensitive to some of the other bile acids tested (Fig. 6A). In contrast, no statistically significant changes in bile acid sensitivity were observed in the Δ*hexSDF* and Δ*ugtB* mutants (Fig. 6A).

**Figure 6.**
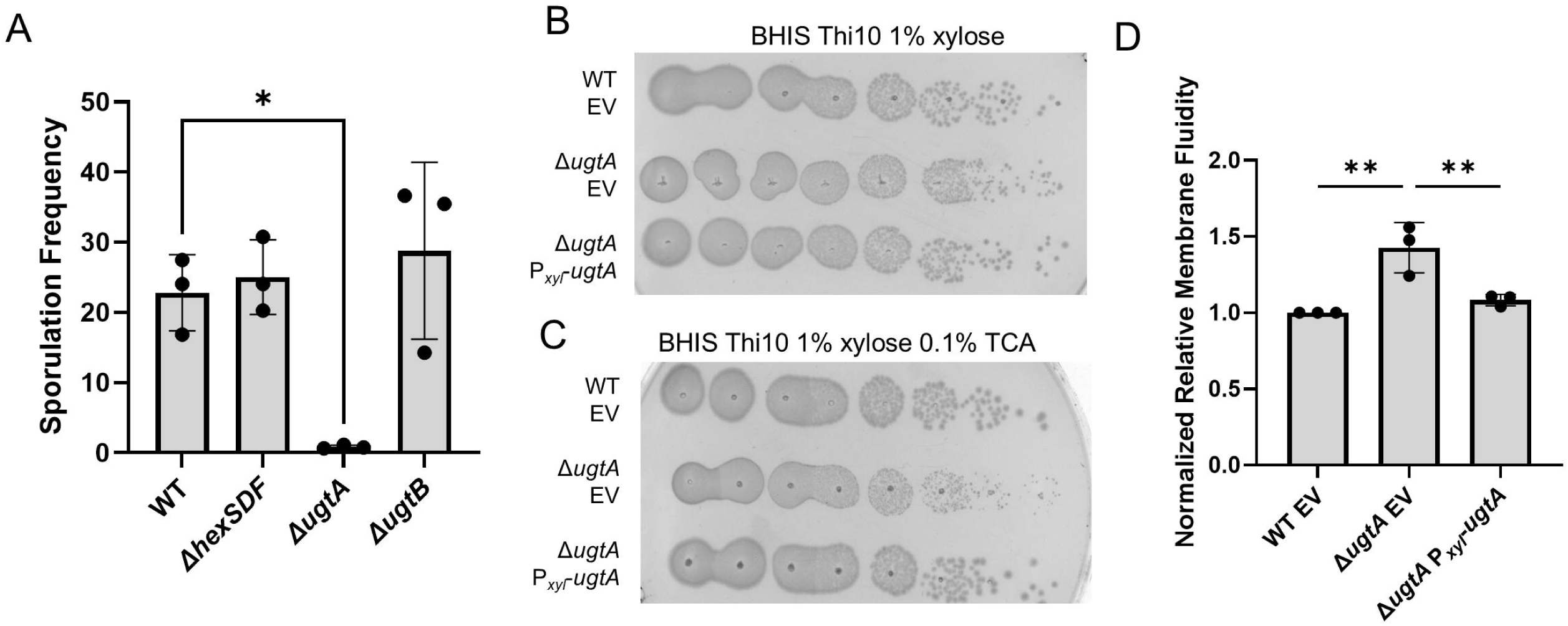
Loss of glycolipids decreases resistance to multiple membrane targeting antimicrobials. Fold change compared to WT for MICs of WT, Δ*hexSDF,* Δ*ugtB,* and Δ*ugtA* for A) TCA, GCA, CA, CDCA, DCA, LCA, B) polymyxin B and surfactin, and C) novobiocin. Data are graphed as the mean and standard deviation of three replicates. Statistical significance was assessed by two-way analysis of variance using Dunnett’s multiple-comparison test. **** *p* < 0.0001, *** *p* < 0.001, ** *p* < 0.01. .

We also tested the glycolipid mutants for sensitivity to multiple cell wall or cell membrane targeting antimicrobials and included novobiocin, a DNA synthesis inhibitor, as a control (Fig. 6C, Table S2). In most cases the Δ*ugtB* and Δ*hexSDF* mutants showed no change in sensitivity (Fig. 6, Table S2). The Δ*ugtA* mutant on the other hand had a slightly lower MIC to a wide range of compounds including the control novobiocin suggesting a potential membrane defect (Fig. 6C, Table S2). The Δ*ugtA* mutant was ∼8 fold more sensitive to polymyxin B and ∼4 fold more sensitive to surfactin both of which target the membrane (Fig. 6B). Expression of *ugtA* in a Δ*ugtA* mutant restored the polymyxin B, surfactin, and novobiocin MIC to levels comparable to wild type with an empty vector control (Fig. S5). Also as previously reported, the Δ*hexSDF* mutant was more sensitive to daptomycin and bacitracin (Table S2) ^23^.

### Loss of UgtA increases membrane fluidity

Since a Δ*ugtA* mutant lacks glycolipids which normally make up ∼50% of the polar lipids in the membrane and we observed increased sensitivity to compounds known to target the membrane, we sought to test if loss of UgtA altered membrane fluidity. To assess relative membrane fluidity, we used the fluorescent dye laurdan which inserts into the membrane and has a shift in emission wavelength depending on the amount of water molecules present in the membrane ^43^. We found that Δ*ugtA* has increased membrane fluidity compared to wild type (Fig. 5D, S3C). When *ugtA* was expressed in the Δ*ugtA* mutant, the relative membrane fluidity returned to levels comparable to a wild-type control (Fig. 5D). In contrast, the Δ*ugtB* and Δ*hexSDF* mutants have the same relative membrane fluidity as wild type (Fig. S3C).

## Discussion

About 50% of the polar lipids of the *C. difficile* membrane are glycolipids, including MHDRG, DHDRG, HNHDRG, and THDRG^20^. This is a higher percentage than in other organisms. Here we identified genes required for synthesis of MHDRG, DHDRG, THDRG, and HNHDRG. We also propose a model which encompasses the synthesis of nearly all the *C. difficile* glycolipids. While the exact roles of glycolipids in the membrane are unknown, they appear to be critical for optimal sporulation, maintenance of cell shape, membrane integrity, and resistance to multiple membrane targeting antimicrobials.

### Model for glycolipid synthesis

Our data show that UgtA is required for synthesis of all glycolipids in *C. difficile* (Fig. 2). UgtB is required for synthesis of DHDRG and THDRG, while HexSDF are required for synthesis of HNHDRG (Fig. 2) ^23^. Based on these data we propose a model where UgtA does not function processively and produces only MHDRG. MHDRG can then be used as a substrate by either HexSDF to produce HNHDRG or UgtB to produce DHDRG and THDRG. This model is supported by data showing that expression of *ugtA* in *B. subtilis* Δ*ugtP* is sufficient for production MHDRG but not the other glycolipids. Conversely, *B. subtilis* Δ*ugtP* expressing *hexSDF* or *ugtB* alone fails to produce any detectable glycolipids (Fig. 3). However, when *hexSDF* were expressed along with *ugtA* in *B. subtilis* Δ*ugtP,* both MHDRG and HNHDRG were produced. Likewise, when *ugtA* and *ugtB* were co-expressed in *B. subtilis* Δ*ugtP* we detected MHDRG and a small amount of DHDRG (Fig. 3). However, it is unclear if UgtB is acting processively to synthesize both DHDRG and THDRG.

The specific sugars that comprise each of the *C. difficile* glycolipids have not been experimentally determined, but the LTA (PS-III) structure has been solved and shows it is anchored by triglucosyldiacylglycerol, which we presume to be THDRG ^15^. While the glycolipids may consist of multiple sugars, it follows that since a triglucosyl glycolipid acts as the LTA anchor, that MHDRG, DHDRG, and THDRG are likely composed of glucose molecules. It is important to note that one cannot assume loss of glycolipids also results in loss of LTA because when the glycolipid anchor is absent in other organisms, LTA is synthesized on another lipid anchor ^27,29,30^. However, future work will be required to determine the effect of loss of glycolipids on LTA biosynthesis.

### Glycolipids are required for optimal sporulation

The ability of *C. difficile* to form spores is critical for its transmission as spores are metabolically dormant cells that can persist in an aerobic environment and resist many antimicrobials. While the membrane lipid composition of the spores of *C. difficile* is not known*, B. subtilis* spores contain glycolipids ^18,19^. Nevertheless, loss of glycolipids in a *B. subtilis ugtP* mutant does not result in a sporulation defect ^18^. In contrast, loss of all glycolipids in *C. difficile* reduced sporulation frequency ∼25-fold as seen with Δ*ugtA* (Fig. 5A). However, loss of DHDRG and THDRG (Δ*ugtB*) or HNHDRG (Δ*hexSDF*) did not alter sporulation frequency (Fig. 5A). Whether the sporulation defect is due to loss of a specific glycolipid or the loss of such a large percentage of the normal polar lipids is unclear. It is also unclear why glycolipids are important for sporulation in *C. difficile* but not *B. subtilis*, but a likely possibility is that glycolipids make up a higher proportion of the polar lipids in *C. difficile* than in *B. subtilis*.

### Glycolipids provide protection against cell membrane targeting antimicrobials

To colonize the host*, C. difficile* spores must germinate into vegetative cells in the intestinal track. The primary germinant signal is the conjugated primary bile acid TCA, but GCA, DCA, and CA can also act as germinant signals ^9^. The secondary bile acids LCA and DCA inhibit *C. difficile* vegetative cells after germination ^9,12,13^. Bile acids are detergents that can exert antimicrobial activity by disrupting bacterial membranes ^44^. *C. difficile* is resistant to conjugated primary bile acids ^13^. We discovered that Δ*ugtA* is significantly more sensitive than wild type to the conjugated primary bile acids TCA and GCA, and only slightly more sensitive to the secondary and unconjugated primary bile acids tested to which *C. difficile* is naturally more sensitive (Fig. 6). This raises the possibility that the high percentage of glycolipids in *C. difficile* is important in mediating resistance to the germinant signals like the conjugated primary bile acid TCA.

We also found that Δ*ugtA* has decreased resistance to polymyxin B and surfactin, which destabilize membranes, and that the Δ*ugtA* mutant membrane is more fluid than the wild type membrane (Fig. 5D) ^45,46^. This increase in fluidity may explain the modest increase in novobiocin sensitivity of the Δ*ugtA* mutant which may be due to a “leaky” membrane. The sensitivities to membrane targeting compounds raise the possibility that the *ΔugtA* mutant may be more sensitive to host defenses, including membrane targeting antimicrobial peptides. It was previously reported that an *S. agalactiae* mutant that lacks glycolipids was more sensitive to killing by neutrophils and cationic antimicrobial peptides and had decreased resistance to some membrane targeting antimicrobials ^29^. This further supports the idea that glycolipids play an important role in maintaining cell membrane integrity and resistance to membrane targeting antimicrobials.

### UgtA is required for normal cell size and morphology

*B. subtilis* UgtP localizes to the site of cell division, where it regulates FtsZ assembly to coordinate cell size with growth rate and nutrient availability ^47^. Loss of UgtP results in *B. subtilis* cells that are shorter and have additional morphological defects ^47,48^. It is possible UgtA in *C. difficile* functions similar to UgtP in *B. subtilis*. This could explain our observation that loss of UgtA results in morphological defects and cells with multiple septa. This might also explain why *C. difficile* Δ*ugtB* is slightly longer than WT because there is likely to be an increase in the UDP-hexose substrate (presumably UDP-glucose) normally used to synthesize DHDRG and THDRG. *C. difficile* might perceive elevated UDP-hexose as a “high nutrient” condition, causing UgtA to inhibit cell division and resulting in longer cells. Alternatively, the absence of all glycolipids in the Δ*ugtA* mutant and the overabundance of MHDRG in the Δ*ugtB* mutant may impair normal cell division by disrupting the physical properties of the cytoplasmic membrane.

## Materials and Methods

### Bacterial strains, media, and growth conditions

Bacterial strains are listed in Table 1. The *C. difficile* strains used in this study are derivatives of R20291. *C. difficile* strains were grown on TY medium consisting of 3% tryptone, 2% yeast extract, and 2% agar (for solid medium). TY was supplemented as needed with thiamphenicol at 10 µg/mL (Thi10). Conjugations were performed on solid brain-heart infusion (BHI) media (3.65% BHI, 2% agar) and plated on TY with Thi10, kanamycin at 50 µg/mL, and cefoxitin at 8 µg/mL. *C. difficile* strains were maintained at 37°C in an anaerobic chamber (Coy Laboratory Products) in an atmosphere of ∼2% H_2_, ∼5% CO_2_, and ∼93% N_2_. *E. coli* strains were grown in lysogeny broth (LB) medium (1% tryptone, 0.5% yeast extract, 0.5% NaCl, and 1.5% agar for solid medium) at 37°C with chloramphenicol at 10 µg/mL (Cam10) and ampicillin at 100 µg/mL (Amp100) as needed. *B. subtilis* strains were grown in LB medium at 37°C with Amp100, spectinomycin 100 µg/mL, or MLS (erythromycin 1 µg/mL plus lincomycin 25 µg/mL) as needed.

**Table 1.**
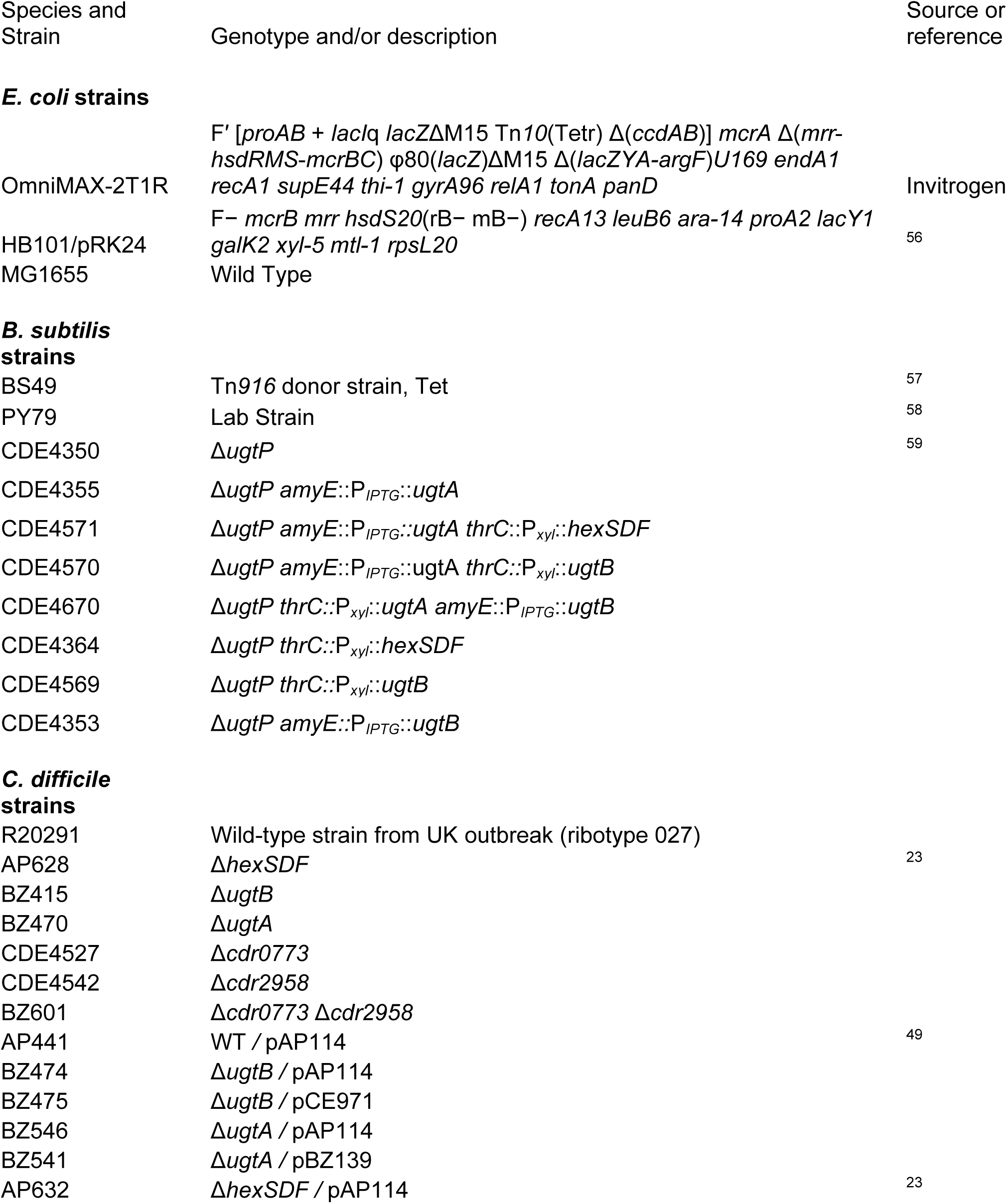
Strains.

### Plasmid and bacterial strain construction

All plasmids are listed in Tables 2 and S3. Plasmids were constructed using isothermal assembly (ITA) (NEBuilder HiFi DNA Assembly, New England Biolabs, Ipswich, MA). Regions of the plasmids constructed using PCR were verified by DNA sequencing. Oligonucleotide primers used in this work were synthesized by Integrated DNA Technologies (Coralville, IA) and are listed in Table S4. All plasmids were propagated using OmniMax-2 T1R as a cloning host (except pCE1069 and pCE1071, which used Able K [Agilent]). For xylose-inducible expression constructs in *C. difficile*, genes of interest were amplified using PCR and inserted into the plasmid pAP114 at the SacI and BamHI sites, as described previously ^49^. CRISPR-Cas9 plasmids were designed as previously described ^50,51^. Plasmids were conjugated into *C. difficile* using either HB101/pRK24 or *B. subtilis* BS49 as previously described ^50–52^.

**Table 2.**
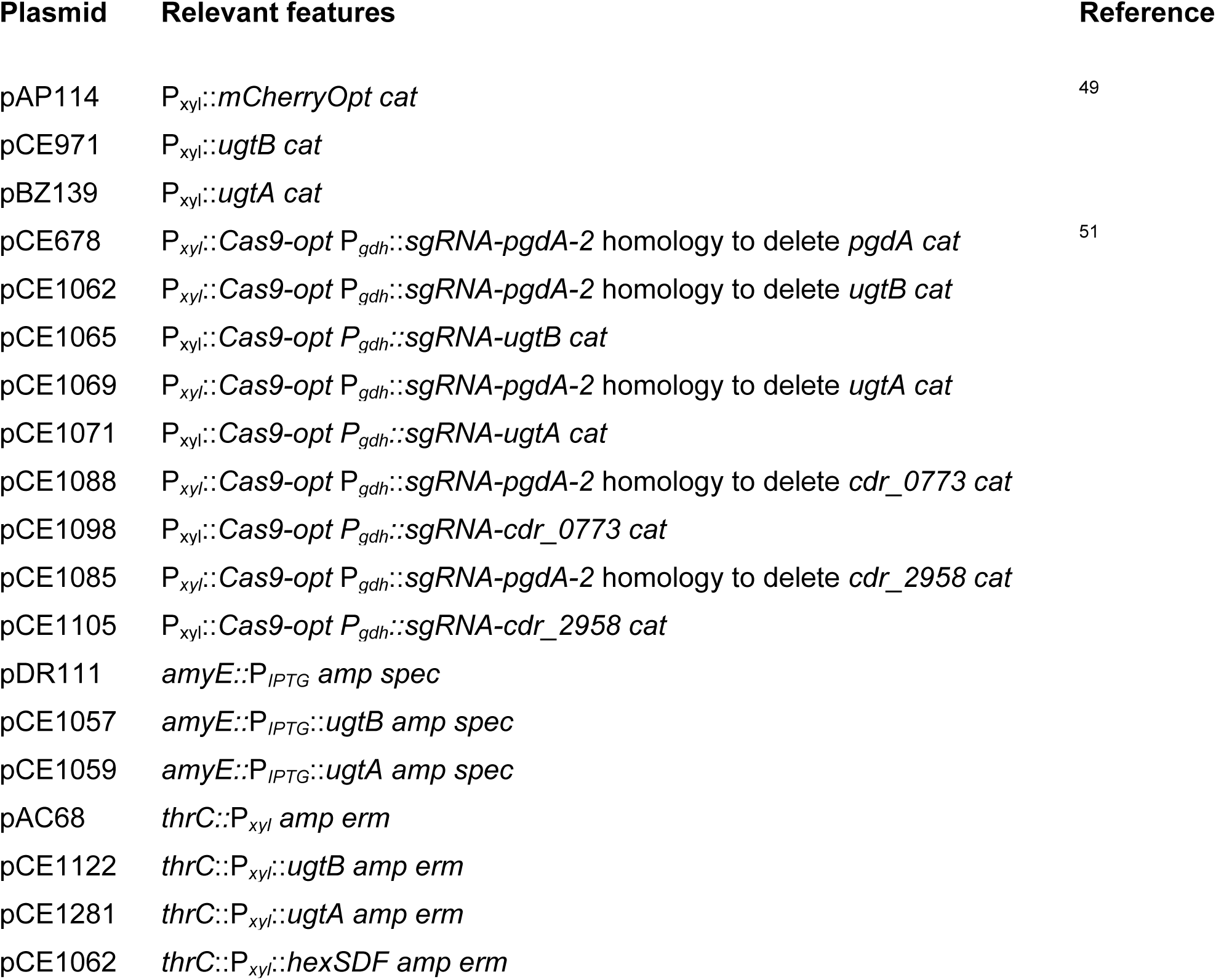
Plasmids.

| Plasmid | Relevant features | Reference |
| --- | --- | --- |
| pAP114 | $P_{xyl}::mCherryOpt\ cat$ | 49 |
| pCE971 | $P_{xyl}::ugtB\ cat$ | |
| pBZ139 | $P_{xyl}::ugtA\ cat$ | |
| pCE678 | $P_{xyl}::Cas9-opt\ P_{gdh}::sgRNA-pgdA-2$ homology to delete *pgdA* *cat* | 51 |
| pCE1062 | $P_{xyl}::Cas9-opt\ P_{gdh}::sgRNA-pgdA-2$ homology to delete *ugtB* *cat* | |
| pCE1065 | $P_{xyl}::Cas9-opt\ P_{gdh}::sgRNA-ugtB\ cat$ | |
| pCE1069 | $P_{xyl}::Cas9-opt\ P_{gdh}::sgRNA-pgdA-2$ homology to delete *ugtA* *cat* | |
| pCE1071 | $P_{xyl}::Cas9-opt\ P_{gdh}::sgRNA-ugtA\ cat$ | |
| pCE1088 | $P_{xyl}::Cas9-opt\ P_{gdh}::sgRNA-pgdA-2$ homology to delete *cdr_0773* *cat* | |
| pCE1098 | $P_{xyl}::Cas9-opt\ P_{gdh}::sgRNA-cdr_0773\ cat$ | |
| pCE1085 | $P_{xyl}::Cas9-opt\ P_{gdh}::sgRNA-pgdA-2$ homology to delete *cdr_2958* *cat* | |
| pCE1105 | $P_{xyl}::Cas9-opt\ P_{gdh}::sgRNA-cdr_2958\ cat$ | |
| pDR111 | *amyE*::$P_{IPTG}$ *amp spec* | |
| pCE1057 | *amyE*::$P_{IPTG}::ugtB$ *amp spec* | |
| pCE1059 | *amyE*::$P_{IPTG}::ugtA$ *amp spec* | |
| pAC68 | *thrC*::$P_{xyl}$ *amp erm* | |
| pCE1122 | *thrC*::$P_{xyl}::ugtB$ *amp erm* | |
| pCE1281 | *thrC*::$P_{xyl}::ugtA$ *amp erm* | |
| pCE1062 | *thrC*::$P_{xyl}::hexSDF$ *amp erm* | |

For *B. subtilis* expression plasmids, genes of interest were amplified using PCR and inserted into the plasmid pAC68 at the HindIII and BamHI sites for xylose-inducible constructs and pDR111 at the HindIII and SphI sites for IPTG-inducible constructs. To construct *B. subtilis* strains, plasmids were transformed into *B. subtilis* PY79 as previously described ^53^.

### Lipid extraction

Lipid extractions were performed as previously described ^40^. Overnight cultures of *C. difficile* were grown in TY (supplemented with 1% xylose and Thi10 when needed) and *B. subtilis* were grown in LB (supplemented with 1% xylose and/or 1 mM IPTG when needed). Overnight cultures were subcultured to an OD_600_ of 0.05 into 500 ml TY (supplemented with 1% xylose and Thi10 when needed) for *C. difficile* and 200 ml LB (supplemented with 1% xylose and/or 1 mM IPTG, when needed) for *B. subtilis* and grown to log phase (OD_600_ 0.6-0.7), then pellets were harvested at 3,000 x *g* for 10 min. Cell pellets were washed in cold 20 mM MOPS, pH 7.2, resuspended in 1 ml water, transferred to a glass centrifuge tube, and 3.75 ml of chloroform-methanol (1:2, vol/vol) were added. The mixture was incubated at room temperature for 2 hours in a fume hood with the cap off and vortexed periodically. The mixture was then centrifuged for 10 min at 3,000 x *g*, and the supernatant decanted into a clean glass centrifuge tube. The remaining pellet was resuspended in 4.75 ml methanol-chloroform-water (2:1:0.8, vol/vol/vol). The mixture was centrifuged for 10 min at 3,000 x *g*, and the supernatant decanted to combine with the first supernatant. To the supernatants, 2.5 ml each of chloroform and water was added and the mixture was centrifuged for 10 min at 3,000 x *g*. The lower, chloroform phase was, transferred to a polypropylene screw cap centrifuge tube (Corning; Corning, NY), and let sit open overnight to evaporate in a 35°C heating block in a fume hood. The resulting lipid extract was dissolved in chloroform.

### TLC and lipid staining

TLC and lipid staining were performed as previously described, with modifications to the solvent condition ^40^. Silica gel aluminum TLC plates were activated by heating for 30 min at 125°C. Once cooled, 0.5 mg of lipid extracts was spotted ∼1 cm from the bottom of the plate. The plate was placed into a TLC developing tank and run until the solvent reached ∼0.5 cm from the top of the plate. The solvent conditions used were chloroform-methanol-ammonium hydroxide-water (7:2.5:0.25:0.25, vol/vol/vol/vol). After drying, the plate was sprayed with 0.5% 1-naphthol in methanol-water (1:1, vol/vol) and then 4.25M H_2_SO_4_ using a glass atomizer. The plate was incubated for 15 min at 125°C to visualize glycolipids.

### Lipidomics

Overnight cultures of *C. difficile* were grown in TY (supplemented with 1% xylose and Thi10 when needed) and *B. subtilis* were grown in LB (supplemented with 1% xylose and/or 1 mM IPTG when needed). Overnight cultures were then subcultured to an OD_600_ of 0.05 into 500 mL TY (supplemented with 1% xylose and Thi10 when needed) for *C. difficile* and 500 mL LB (supplemented with 1% xylose and/or 1 mM IPTG when needed) for *B. subtilis*. Subcultures were allowed to grow until an OD_600_ of 0.6 to 0.7 was reached, at which point the cells were harvested and pelleted at 3,000 × g for 10 min. Biological replicates were grown on different days. Lipidomics were performed in biological triplicate for *C. difficile* wild type, Δ*ugtA,* and Δ*ugtB*. For the *C. difficile* complementation and *B. subtilis* strains, a single biological replicate was performed as we were only interested in the detection of the presence of lipids species, not the formal quantitative comparison as there are artificial levels of expression of elements of interest in these strains.

Lipid extraction and liquid chromatography with tandem mass spectrometry (LC-MS/MS) were performed by Cayman Chemical Company. After thawing, cells were mixed with 5 mL methanol, transferred to 7 mL Precellys tubes containing 0.1 mm ceramic beads (Bertin Technologies; CK01 lysing kit), and homogenized with three cycles at 8,800 rpm for 30 s, with 60 s pauses between cycles. Then, 800 μL of the homogenized mixtures was transferred to 8 mL screw-cap glass tubes. A methyl tert-butyl ether (MTBE)-based liquid-liquid extraction protocol was used by first adding 1.2 mL methanol containing a mixture of deuterated internal standards covering several major lipid categories (fatty acids, glycerolipids, glycerophospholipids, sphingolipids, and sterols) and then 4 mL MTBE. The mixture was incubated on a tabletop shaker at 500 rpm at room temperature for 1 hour and then stored at 4°C for 60 hours to maximize lipid extraction. After bringing the samples to room temperature, phase separation was induced by adding 1 mL water to each sample. The samples were vortexed and then centrifuged at 2,000 × *g* for 15 min. The upper organic phase of each sample was carefully removed using a Pasteur pipette and transferred into a clean glass tube. The remaining aqueous phase was reextracted with 2 mL of the upper phase of MTBE/methanol/water at 10:3:2.5 (vol/vol/vol). After vortexing and centrifuging, the organic phase was collected and combined with the initial organic phase. The extracted lipids were dried overnight in a SpeedVac vacuum concentrator.

The dried lipid extracts were reconstituted in 200 μL n-butanol–methanol at 1:1 (vol/vol) and transferred into autosampler vials for analysis by LC-MS/MS. Aliquots of 5 mL were injected into an Ultimate 3000 ultraperformance liquid chromatography system connected to a Q Exactive Plus Orbitrap mass spectrometer (Thermo Scientific). An Accucore C30 2.6 mm, 150 by 2.1 mm HPLC column (Thermo Scientific) was used, using mobile phases A [acetonitrile/water/formic acid 60:40:0.1 (vol/vol/vol), containing 10 mM ammonium formate] and B [acetonitrile/isopropanol/formic acid 10:90:0.1 (vol/vol/vol), containing 10 mM ammonium formate]. Lipids were eluted at a constant flow rate of 300 mL/min using a gradient from 30% to 99% mobile phase B over 30 min. The column temperature was kept at a constant 40°C. Polarity switching was used throughout the gradient to acquire high-resolution MS data (resolution, 75,000) and data-dependent MS/MS data.

Data analysis was performed using Lipostar software (version 2; Molecular Discovery) for detection of features (peaks with unique m/z and retention time), noise and artifact reduction, alignment, normalization, and lipid identification. Automated lipid identification was performed by querying the Lipid Maps Structural Database (LMSD), modified by Cayman to include many additional lipids not present in the LMSD. To allow for comparison between strains, the summed peak areas of lipids with the same head groups were normalized to the wild-type values.

### Microscopy

Overnight cultures of *C. difficile* were subcultured in TY to an OD_600_ of 0.05 (supplemented with Thi10 and 1% xylose when needed) and allowed to grow until an OD_600_ of 0.6 to 0.7. Cells were immobilized using thin agarose pads (1%). Phase-contrast micrographs were recorded on an Olympus BX60 microscope equipped with a 100× UPlanApo objective (numerical aperture, 1.35). Micrographs were captured with a Hamamatsu Orca Flash 4.0 V2 + complementary metal oxide semiconductor camera. Excitation light was generated with an X-Cite XYLIS LED light source. Membranes were stained with the lipophilic dye FM4-64 (Life Technologies) at 10 µg/mL. Cells were imaged immediately without washing. Red fluorescence was detected with the Chroma 49008 filter set (538 to 582 nm excitation filter, 587 nm dichroic mirror, and a 590 to 667 nm emission filter). The plug-in module MicrobeJ from the image analysis package Fiji was used to measure cell length and sinuosity ^54,55^. Cell sinuosity is the ratio of the cell length along its medial axis and the distance between the poles of a cell. A cell with a sinuosity value of 1 is a perfectly straight rod while curvy cells have sinuosity values larger than 1. At least 300 cells from three independent experiments were used for quantification.

### Sporulation

Sporulation frequencies were determined as previously described ^42^. Briefly, overnight cultures were grown and subcultured in BHIS (3.7% BHI, 0.5% yeast extract) with 0.1% TCA and 0.2% fructose, plated onto 70:30 sporulation agar (6.3% bacto peptone, 0.35% protease peptone, 1.11% BHI, 0.15% yeast extract, 0.106% tris base, 0.07% ammonium sulfate, 1.5% agar), and grown for 24 hours at 37°C. A cell suspension was made with cells scraped off the 70:30 media plates, and a portion of the cell suspension was treated with 28.5% ethanol (final concentration) to kill vegetative cells. Dilutions of the untreated cell suspension were plated onto BHIS and CFU/ml of vegetative cells was calculated from colony counts after incubation for 24 hours at 37°C. Dilutions of the ethanol killed cells were plated onto BHIS 0.1% TCA and CFU/ml of spores was calculated from colony counts after incubation for 24 hours at 37°C. Sporulation frequency was calculated using the following formula: [total spore CFU/ml / total cell CFU/ml (vegetative cells CFU/ml + spores CFU/ml)] *100. For strains carrying plasmids, Thi10 and 1% xylose were added to overnight cultures, subcultures, and 70:30 plates. Data are represented as an average from three independent experiments.

### Membrane fluidity

Laurdan (6-Dodecanoyl-N,N-dimethyl-2-naphthylamine) (Sigma-Aldrich, catalog number: 40227) was used to assess relative membrane fluidity as previously described ^43^. Briefly, overnight cultures of *C. difficile* were subcultured to an initial OD_600_ of 0.05 and grown to an OD_600_ 0.6-0.7. Cells were treated with 10 μM laurdan for 10 min in the dark at 37°C in an anaerobic chamber. Cells were then removed from the anaerobic chamber and washed four times with pre-warmed laurdan buffer (137 mM NaCl, 2.7 mM KCl, 10 mM Na_2_HPO_4_, 1.8 mM KH_2_PO_4_, 0.2% glucose, 1% DMF) and resuspended to a final OD_600_ of 0.5 in pre-warmed laurdan buffer. Cell suspensions were added to a pre-warmed black, flat bottomed 96-well plate and fluorescence was read every min for 20 min with excitation at 350 nm and emission at 460 nm and 500 nm. Relative membrane fluidity was calculated by (I_460_+I_500_) / (I_460_-I_500_) at the five-minute time point and normalized to wild type. For strains carrying plasmids, Thi10 and 1% xylose were added to all cultures. Data represented are an average of three independent experiments.

### Antimicrobial MIC determination

Overnight cultures of *C. difficile* were subcultured, grown to late log phase (OD_600_ 1.0), and then diluted into TY to 10^6^ CFU/ml. A series of antibiotic concentrations was prepared in a 96-well plate in 50 μl of TY. Wells were inoculated with 50 μl of the dilute late-log phase culture (0.5 × 10^5^ CFU/well) and grown at 37°C for 16 hours. After incubation, the MIC was determined based on the presence of cell pellets. For lysozyme MICs, 10 μl from each well was diluted 1:10 in TY broth and 5 μl was plated onto TY agar and incubated for 24 hours. The MIC was determined based on the lowest concentration of lysozyme where 5 or fewer colonies were found per spot. Data are reported as the average from three independent experiments.

### Transmission Electron Microscopy (TEM)

Overnight cultures of *C. difficile* were subcultured and grown to log phase (OD_600_ 0.6-0.7) in Thi10 and 1% xylose and fixed with 2.5 % glutaraldehyde (in 0.1 M sodium cacodylate buffer, pH 7.4) overnight at 4°C. Samples were postfixed with 1% osmium tetroxide for 1 hr and then rinsed in 0.1 M sodium cacodylate buffer. Following serial alcohol dehydration (50%, 75%, 95%, 100%), the samples were embedded in Epon 12 (Ted Pella, Redding, CA). Ultramicrotomy was performed, and ultrathin sections (70 nm) were poststained with uranyl acetate and lead citrate. Samples were examined with a Hitachi HT-7800 transmission electron microscope (TEM) (Tokyo, Japan).

## Acknowledgements

This work was supported by Public Health Service Grant R01AI087834 to C.D.E. from the National Institute of Allergy and Infectious Diseases. B.R.Z. was supported by grant T32GM008365 and the University of Iowa Center for Biocatalysis and Bioprocessing. The *ugtP* mutant was provided by the Bacillus Genetic Stock Center (BGSC). The TEM images presented here were obtained at the University of Iowa Central Microscopy Research Facility, a core resource supported by the University of Iowa Vice President for Research, and the Carver College of Medicine. We thank members of the Ellermeier and Weiss laboratories for helpful discussions.

