## Supplementary figures and images for "Identification of two glycosyltransferases required for synthesis of membrane glycolipids in *Clostridioides difficile*"

### Fig. S1

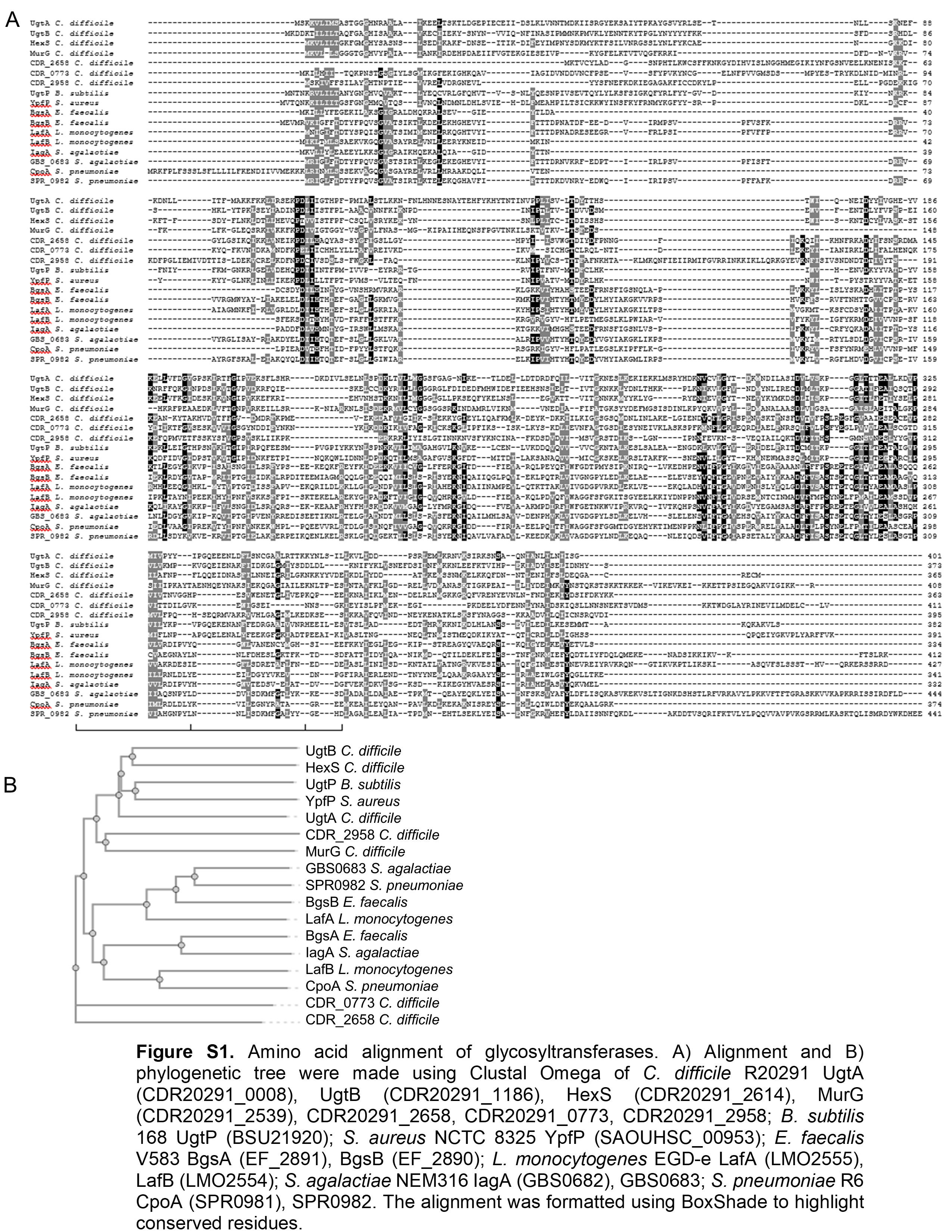

### Fig. S2

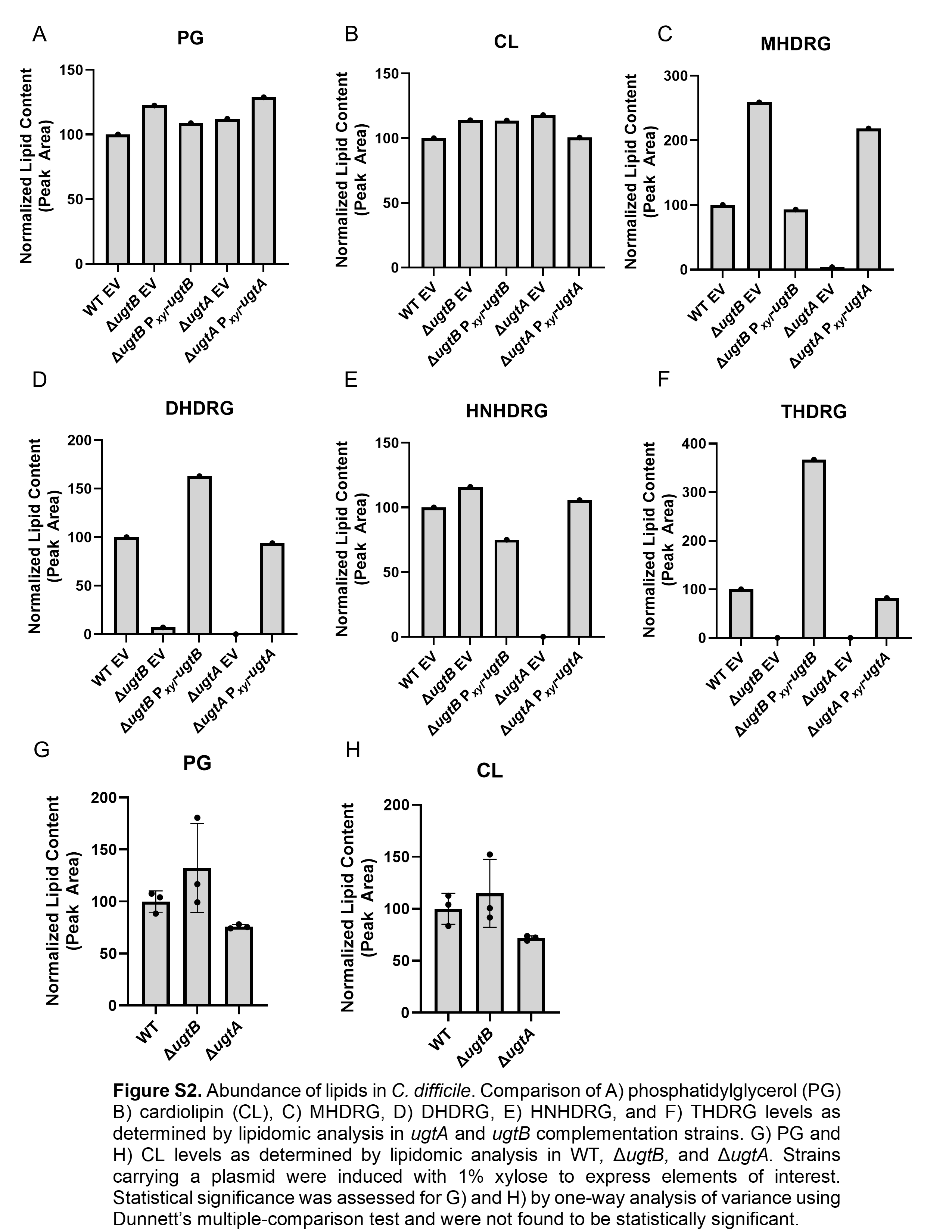

### Fig. S3

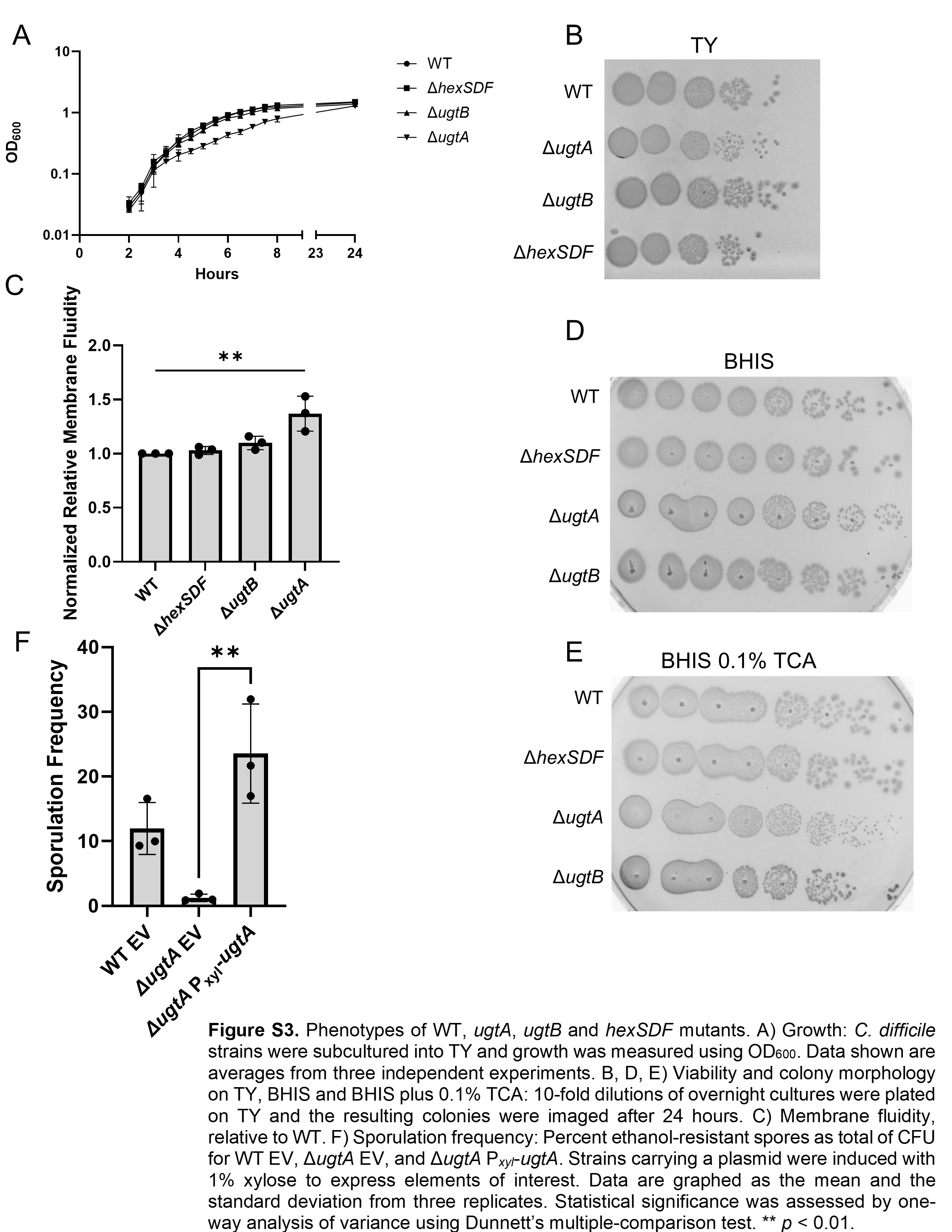

### Fig. S4

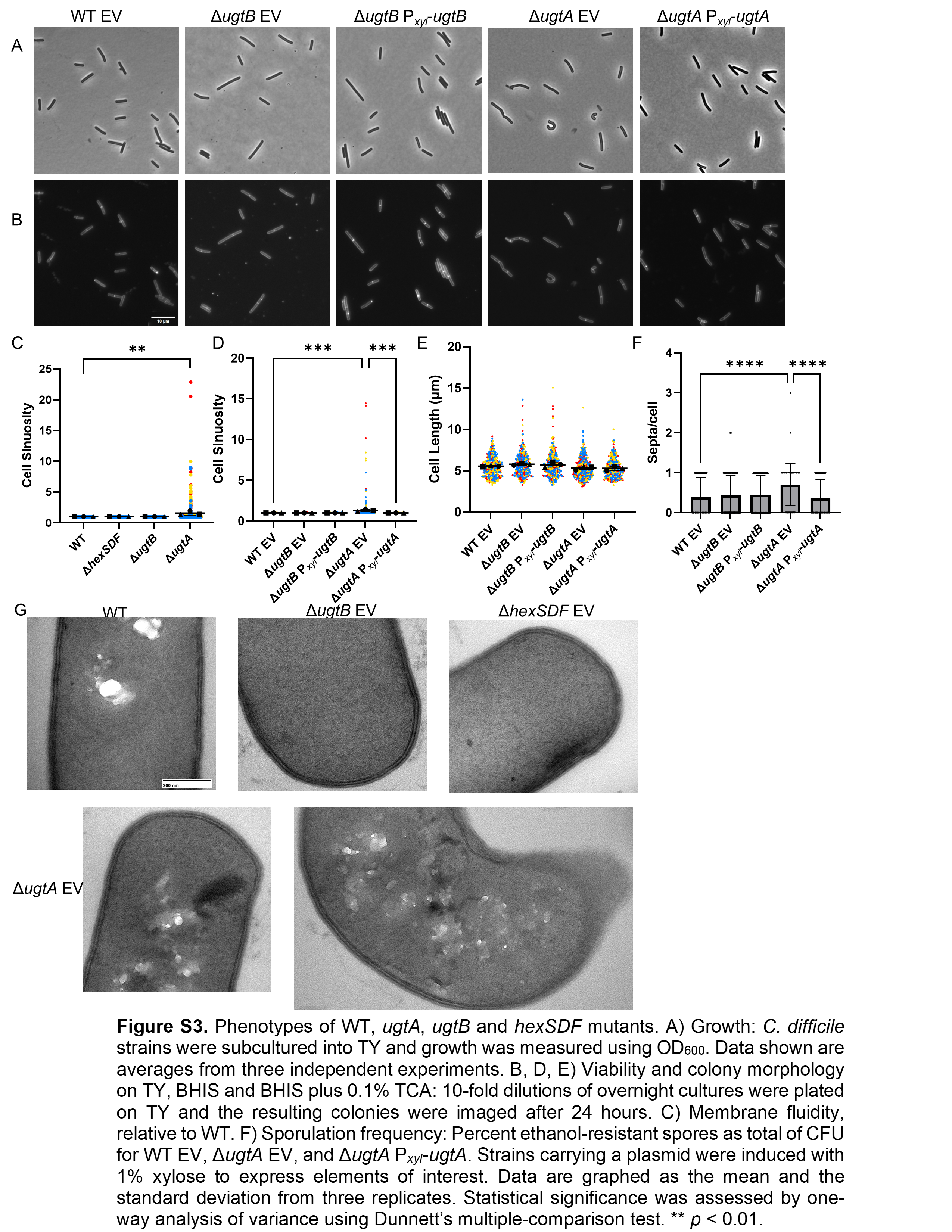

### Fig. S5

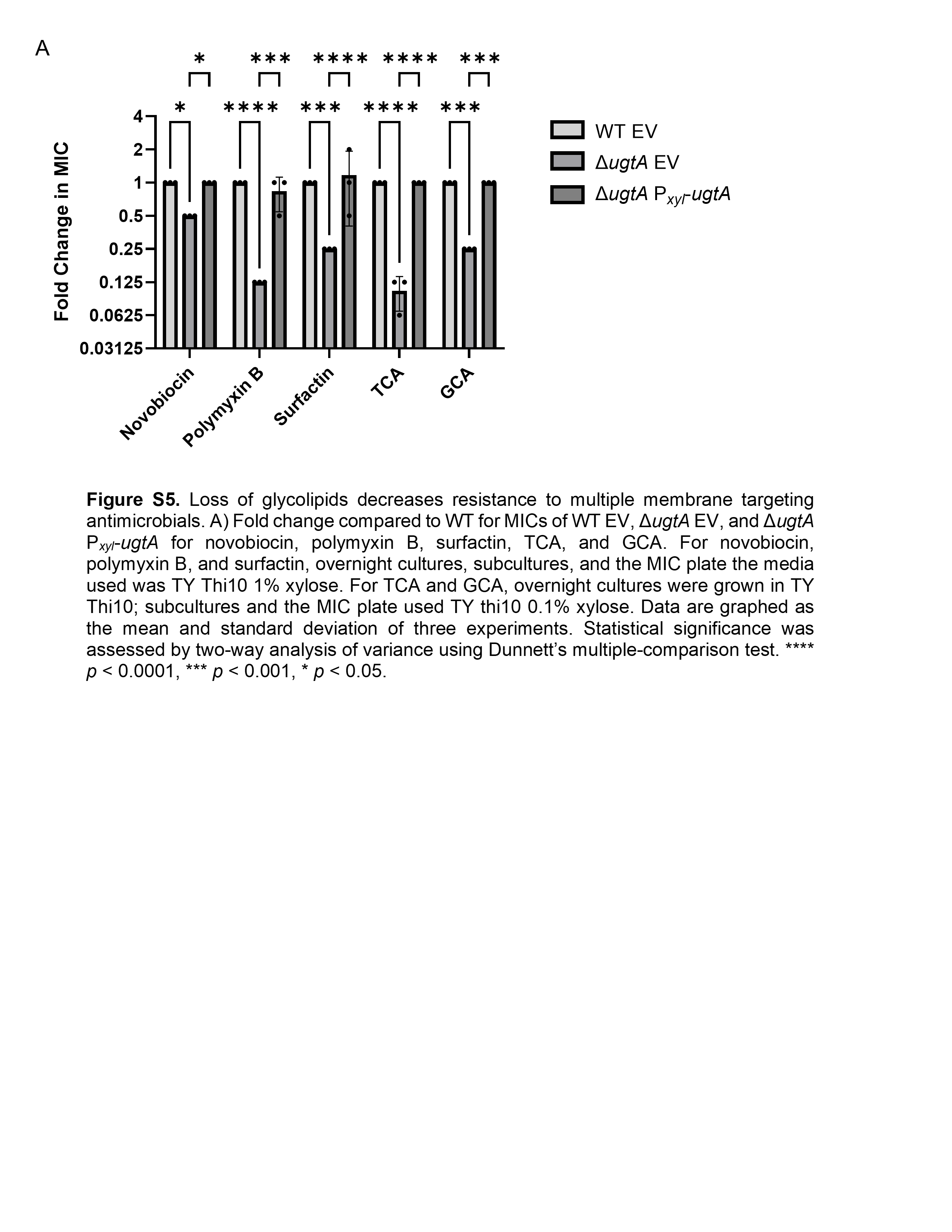
