## Supplementary material for "Identification of two glycosyltransferases required for synthesis of membrane glycolipids in *Clostridioides difficile*": Table S1

**Table S1 BLAST Results of diacylglycerol glucosyltransferases**

| Protein | Product Name | Score (Bits) | E Value | Query Protein | Query Organism | PFAM Family | PFAM Description |
|---|---|---|---|---|---|---|---|
| CDR20291_0008 | putative glycosyl transferase | 164 | 5.00E-48 | UgtP | *B. subtilis* | PF06925.16;PF04101.21 | MGDG_synth;Glyco_tran_28_C |
| CDR20291_1186 | putative cell wall biosynthesis protein | 146 | 3.00E-41 | UgtP | *B. subtilis* | PF06925.16;PF04101.21 | MGDG_synth;Glyco_tran_28_C |
| HexS | putative monogalactosyldiacylglycerol synthase | 129 | 4.00E-35 | UgtP | *B. subtilis* | PF06925.16;PF04101.21 | MGDG_synth;Glyco_tran_28_C |
| CDR20291_0773 | putative glycosyl transferase | 58.5 | 1.00E-10 | LafB | *L. monocytogenes* | PF13439.11;PF00534.25 | Glyco_transf_4;Glycos_transf_1 |
| CDR20291_2658 | putative capsular polysaccharide biosynthesis glycosyl transferase | 48.5 | 4.00E-07 | SPR_0982 | *S. pneumoniae* | PF13477.11;PF00534.25 | Glyco_trans_4_2;Glycos_transf_1 |
| MurG | UDP-N-acetylglucosamine--N-acetylmuramyl-(penta peptide) pyrophosphoryl-undecaprenol N-acetylglucosamine transferase | 47.4 | 8.00E-07 | UgtP | *B. subtilis* | PF03033.25;PF04101.21 | Glyco_transf_28;Glyco_tran_28_C |
| CDR20291_2958 | putative glycosyltransferase | 43.1 | 2.00E-05 | UgtP | *B. subtilis* | PF00201.23 | UDPGT |
