## Supplementary material for "Identification of two glycosyltransferases required for synthesis of membrane glycolipids in *Clostridioides difficile*": Table S2

**Table S2 MIC**

| **MIC** | WT | Δ*hexSDF* | Δ*ugtB* | Δ*ugtA* |
|---|---|---|---|---|
| Ampicillin (µg/ml) | 1.33 ± 0.58 | 0.8 ± 0 | 1 ± 0 | 0.75 ± 0.43 |
| Bacitracin (µg/ml) | 256 ± 0 ^ | 64 ± 0 ^ | 256 ± 0 ^ | 128 ± 0 ^ |
| Cefoxitin (µg/ml) | 256 ± 0 ^ | ND | ND | 256 ± 0 ^ |
| Daptomycin (µg/ml) | 4 ± 0 | **0.833 ± 0.29 ****** | 4 ± 0 | 4 ± 0 |
| Fosfomycin (µg/ml) | 16 ± 0 ^ | ND | 16 ± 0 ^ | 32 ± 0 ^ |
| Meropenem (µg/ml) | 3.33 ± 1.15 | ND | ND | 3.33 ± 1.15 |
| Nisin (µg/ml) | 180 ± 0 | 120 ± 51.96 | **90 ± 0 **** | **90 ± 0 **** |
| Vancomycin (µg/ml) | 1 ± 0 ^ | ND | 0.5 ± 0 ^ | 1 ± 0 ^ |
| Lauric Acid (mM) | 0.25 ± 0 ^ | 0.25 ± 0 ^ | 0.25 ± 0 ^ | 0.25 ± 0 ^ |
| Linoleic Acid (mM) | >32 | >32 | >32 | >32 |
| Lysozyme (mg/ml) | 8 ± 0 | 6.67 ± 2.31 | **4 ± 0 *** | **3.33 ± 1.15 **** |
| Novobiocin (µg/ml) | 16 ± 0 | 16 ± 0 | 16 ± 0 | 10.67 ± 4.62 |

ND-not determined
Mean ± standard deviation
* $p < 0.05$
** $p < 0.01$
**** $p < 0.0001$
^Did not calculate statistical significance using one-way analysis of variance as the standard deviation was 0.
