## Supplementary material for "Identification of two glycosyltransferases required for synthesis of membrane glycolipids in *Clostridioides difficile*": Table S3

**Table S3 Plasmids**

| Plasmid | Relevant features | Parent vector | Restriction enzymes to digest parent plasmid | Primers used | PCR template | Assembly | Reference |
|---|---|---|---|---|---|---|---|
| pAP114 | $P_{xyl}$::*mCherryOpt cat* | | | | | | 50 |
| pCE971 | $P_{xyl}$::*ugtB cat* | pAP114 | BamHI, SacI | CDEP5693-5694 | R20291 | ITA | |
| pBZ139 | $P_{xyl}$::*ugtA cat* | pAP114 | BamHI, SacI | CDEP6171-6172 | R20291 | ITA | |
| pCE678 | $P_{xyl}$::*Cas9-opt* $P_{gdh}$::*sgRNA-pgdA-2* homology to delete *pgdA cat* | | | | | | 52 |
| pCE1062 | $P_{xyl}$::*Cas9-opt* $P_{gdh}$::*sgRNA-pgdA-2* homology to delete *ugtB cat* | pCE678 | NotI, XhoI | CDEP6257-6260 | R20291 | ITA | |
| pCE1065 | $P_{xyl}$::*Cas9-opt* $P_{gdh}$::*sgRNA-ugtB cat* | pCE1062 | MscI, MluI | CDEP4237, 6271 | pCE678 | ITA | |
| pCE1069 | $P_{xyl}$::*Cas9-opt* $P_{gdh}$::*sgRNA-pgdA-2* homology to delete *ugtA cat* | pCE678 | NotI, XhoI | CDEP6290-6293 | R20291 | ITA | |
| pCE1071 | $P_{xyl}$::*Cas9-opt* $P_{gdh}$::*sgRNA-ugtA cat* | pCE1069 | MscI, MluI | CDEP4237, 6274 | pCE678 | ITA | |
| pCE1088 | $P_{xyl}$::*Cas9-opt* $P_{gdh}$::*sgRNA-pgdA-2* homology to delete *cdr_0773 cat* | pCE678 | NotI, XhoI | CDEP6458-6461 | R20291 | ITA | |
| pCE1098 | $P_{xyl}$::*Cas9-opt* $P_{gdh}$::*sgRNA-cdr_0773 cat* | pCE1088 | MscI, MluI | CDEP4237, 6462 | pCE678 | ITA | |
| pCE1085 | $P_{xyl}$::*Cas9-opt* $P_{gdh}$::*sgRNA-pgdA-2* homology to delete *cdr_2958 cat* | pCE678 | NotI, XhoI | CDEP6454-6457 | R20291 | ITA | |
| pCE1105 | $P_{xyl}$::*Cas9-opt* $P_{gdh}$::*sgRNA-cdr_2958 cat* | pCE1085 | MscI, MluI | CDEP4237, 6469 | pCE678 | ITA | |
| pDR111 | *amyE*:: $P_{IPTG}$ *amp spec* | | | | | | |
| pCE1057 | *amyE*:: $P_{IPTG}$::*ugtB amp spec* | pDR111 | HindIII, SphI | CDEP6229-6230 | R20291 | ITA | |
| pCE1059 | *amyE*:: $P_{IPTG}$::*ugtA amp spec* | pDR111 | HindIII, SphI | CDEP6227-6228 | R20291 | ITA | |
| pAC68 | *thrC*:: $P_{xyl}$ *amp erm* | | | | | | |
| pCE1122 | *thrC*::$P_{xyl}$::*ugtB amp erm* | pAC68 | HindIII, BamHI | CDEP6615-6616 | R20291 | ITA | |
| pCE1281 | *thrC*::$P_{xyl}$::*ugtA amp erm* | pAC68 | HindIII, BamHI | CDEP6628-6629 | R20291 | ITA | |
| pCE1062 | *thrC*::$P_{xyl}$::*hexSDF amp erm* | pAC68 | HindIII, BamHI | CDEP6215-6216 | R20291 | ITA | |
