## Supplementary material for "Identification of two glycosyltransferases required for synthesis of membrane glycolipids in *Clostridioides difficile*": Table S4

**Table S4 Oligos**

| Oligo | Sequence | Relevant Features |
|---|---|---|
| CDEP5693 | cgatagttatgaagtgagcttaaggaggtggctaaaatgaaagatgataaaacaat | Clone *ugtB* onto pAP114 |
| CDEP5694 | gttttattaaaacttataggatcttaactgtaatgattatctattaattctgatatataatcagca | Clone *ugtB* onto pAP114 |
| CDEP6171 | cgatagttatgaagtgagcttaaggaggattttattatgagcaaaaaagtattaataatg | Clone *ugtA* onto pAP114 |
| CDEP6172 | aaagttttattaaaacttataggatcgggattttatttattttatcaaccaga | Clone *ugtA* onto pAP114 |
| CDEP6257 | aaacagctatgaccgcggccccctcatcttcacttattaaaatct | Clone homology for *ugtB* deletion |
| CDEP6258 | tgtttgaaatttttaatagtacaattttagccaatttatcaccactt | Clone homology for *ugtB* deletion |
| CDEP6259 | agtggtgataaattggctaaaattgtactattaaaaatttcaaacataaaatt | Clone homology for *ugtB* deletion |
| CDEP6260 | ttatttttatgctagctcgagaaaaacttagaaatatatgcttcaaaa | Clone homology for *ugtB* deletion |
| CDEP4237 | cttataggatccgcggccgctag | Clone *gRNA* from pCE678 |
| CDEP6271 | aattaaactgtaaatggccactaatgctaactaagcctgggttttagagctagaaatagc | *sgRNA-ugtB* |
| CDEP6290 | aaacagctatgaccgcggccgtccgccgctctttaccgaa | Clone homology for *ugtA* deletion |
| CDEP6291 | ccctaatagataatatatatactatagggcctcctaaaatgtagcatagtatttatattg | Clone homology for *ugtA* deletion |
| CDEP6292 | ctatgctacattttaggaggccctatagtatatatattatctattagggattt | Clone homology for *ugtA* deletion |
| CDEP6293 | ttatttttatgctagctcgagaaataataatggcaacaaatcct | Clone homology for *ugtA* deletion |
| CDEP6274 | aattaaactgtaaatggccagtattagtaagcaaacctgggttttagagctagaaatagc | *sgRNA-ugtA* |
| CDEP6458 | aaacagctatgaccgcggccggagattgtgctttgtttattaaa | Clone homology for *cdr0773* deletion |
| CDEP6459 | agttacactttttataatttaaataatttttcacttgcctcctttctaaaa | Clone homology for *cdr0773* deletion |
| CDEP6460 | agaaaggaggcaagtgaaaaaattatttaaattataaaaaagtgtaactatttttaaac | Clone homology for *cdr0773* deletion |
| CDEP6461 | ttatttttatgctagctcgacttgtatctactctatctacacca | Clone homology for *cdr0773* deletion |
| CDEP6462 | aattaaactgtaaatggccaggataaaaacttacttcactgttttagagctagaaatagc | *sgRNA-cdr0773* |
| CDEP6454 | aaacagctatgaccgcggccctgacagcaccgttattaaa | Clone homology for *cdr2958* deletion |
| CDEP6455 | tgcatgtatgtatatatatttttaacaaatctaatttcatacctccttttataattttttaaaatca | Clone homology for *cdr2958* deletion |
| CDEP6456 | ttataaaaggaggtatgaaattagatttgttaaaaatatatatacatacatgcaatttaatcaa | Clone homology for *cdr2958* deletion |
| CDEP6457 | ttatttttatgctagctcgacctcattacttcttaaaagtttattattattt | Clone homology for *cdr2958* deletion |
| CDEP6469 | aattaaactgtaaatggccaattctctgctccgaatgttggttttagagctagaaatagc | *sgRNA-cdr2958* |
| CDEP6615 | gacctgcaggcatgcaagcttaaggaggtggctaaaatgaa | Clone *ugtB* onto pAC68 |
| CDEP6616 | aaaactgctgccttcggatcttaactgtaatgattatctattaattctgatat | Clone *ugtB* onto pAC68 |
| CDEP6229 | aacaattaagcttaaggaggtggctaaaatgaaagatgataaaac | Clone *ugtB* onto pDR111 |
| CDEP6230 | ccaccgaattagcttgcatgttaactgtaatgattatctattaattctgatat | Clone *ugtB* onto pDR111 |
| CDEP6628 | gacctgcaggcatgcaagctttaggaggattttattatgagcaaa | Clone *ugtA* onto pAC68 |
| CDEP6629 | aaaactgctgccttcggatctcaaccagaaattatatttaaaattaaattagc | Clone *ugtA* onto pAC68 |
| CDEP6227 | aacaattaagcttaaggaggattttattatgagcaaaaaagtattaataatgt | Clone *ugtA* onto pDR111 |
| CDEP6228 | ccaccgaattagcttgcatggggattttatttattttatcaaccaga | Clone *ugtA* onto pDR111 |
| CDEP6215 | ccaccgaattagcttgcatgttatcttacataagcaactttttca | Clone *hexSDF* onto pDR111 |
| CDEP6216 | gagcggataacaattaagctgctttaacgaggagggaatc | Clone *hexSDF* onto pDR111 |
